# Self-assembling protein materials with genetically programmable morphology and size

**DOI:** 10.1101/2025.07.24.666636

**Authors:** Jacob B. Miller, Charlotte H. Abrahamson, Marilyn S. Lee, Brett J. Palmero, Nolan W. Kennedy, Carolyn E. Mills, Danielle Tullman-Ercek

## Abstract

Materials are challenging to synthetically program down to the atom level. Nature, however, excels at creating hierarchical materials from nanoscale building blocks, a feat that remains a major challenge in synthetic systems. A deeper understanding of the molecular rules governing self-assembly would unlock the potential for designing genetically programmable materials with atomic precision. Hexameric bacterial microcompartment (BMC-H) proteins offer a powerful model system for exploring this question. These sequence-defined proteins naturally assemble into complex architectures and can be expressed biologically, making them ideal candidates for studying how minor sequence variations influence supramolecular structure. In this work, we leverage cell-free protein synthesis (CFPS) alongside immunostaining and super-resolution microscopy to investigate the self-assembly behavior of two BMC-H proteins, PduA and PduJ. We find that both proteins form micro-to millimeter scale structures when expressed *in vitro*. Further, we demonstrate how single point mutation changes lead PduA and PduJ to form significantly different supramolecular structures when produced using CFPS. These studies support the future exploration of self-assembling proteins as programmable scaffolds in broad materials applications.

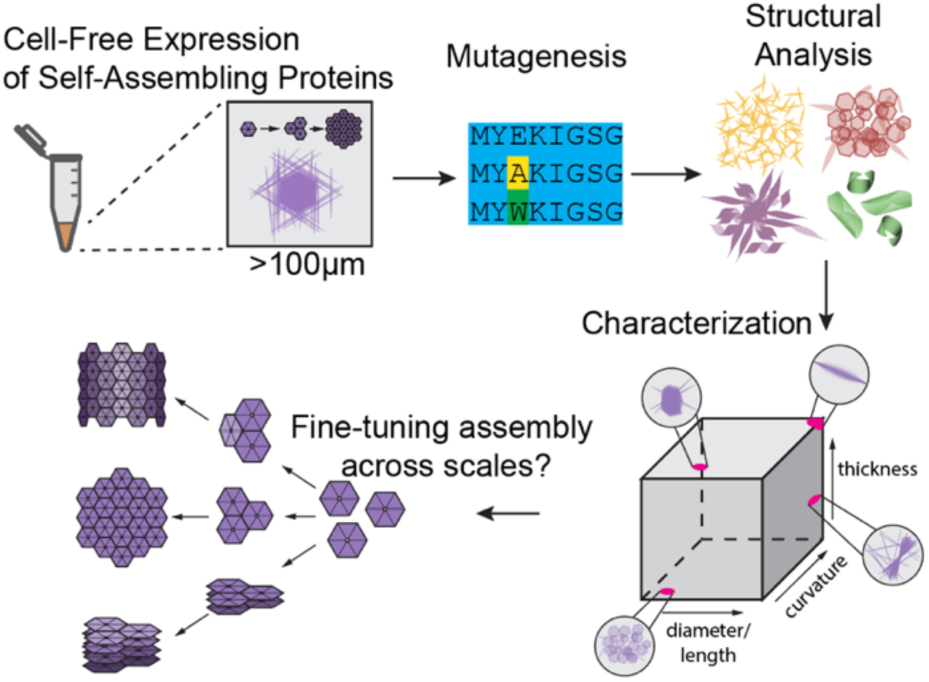

## Introduction

Self-assembling proteins are universal across all kingdoms of life, including viruses, bacteria, fungi, plants, and animals^1^. These proteins are utilized by organisms for diverse functions, from structural elements that impart essential mechanical properties to our tissues^2^, to scaffolds that facilitate intracellular transport^3^, and even as membranes that compartmentalize metabolic processes^4^. Self-assembling proteins are popular candidates for the production and engineering of biomaterials because their genetically encoded nature permits precise control over their sequence and structure. This, in turn, directly controls the interactions governing their assembly and thus dictates the resulting assembled structures^3,5,6^. Because of their ability to adopt a host of different geometries, protein-based materials are already being used for myriad applications, including nanocages for targeted drug delivery^7^, nanotubes for biosensors^8^, and nanosheets for enzyme co-immobilization^9,10^.

<u>B</u>acterial <u>M</u>icro<u>C</u>ompartment – <u>H</u>examer (BMC-H) proteins are one such example of self-assembling proteins found in nature. BMC-H proteins are homohexameric proteins that natively form the structural “shell” of bacterial <u>m</u>icro<u>c</u>om<u>p</u>artments (MCPs)^4^. BMC-H assembly begins when 6 BMC-H proteins monomers hexamerize with circular symmetry (Fig. 1A). Hexamers then tile together through interactions at the interface between two hexamer edges (herein: edge-edge interactions, Fig. 1A)^11^. The angle at which two hexamers associate (herein: edge-edge interaction angle) also dictates the shape of the self-assembled structure^12–15^. In the 1,2-propanediol utilization (Pdu) MCP from *Salmonella enterica* serovar LT2, BMC proteins tile at 0 degrees to form planar facets of the MCP, whereas BMC proteins likely meet at 20 to 30 degree angles near the MCP vertices^13^. In a Pdu MCP homolog from myxobacterium *Haliangium ochraceum,* 25- and 30-degree angles at various vertices between BMC-H proteins regulate the MCP’s icosahedral shape. Moreover, changing these angles leads to formation of structures with different morphologies.

**Figure 1.**
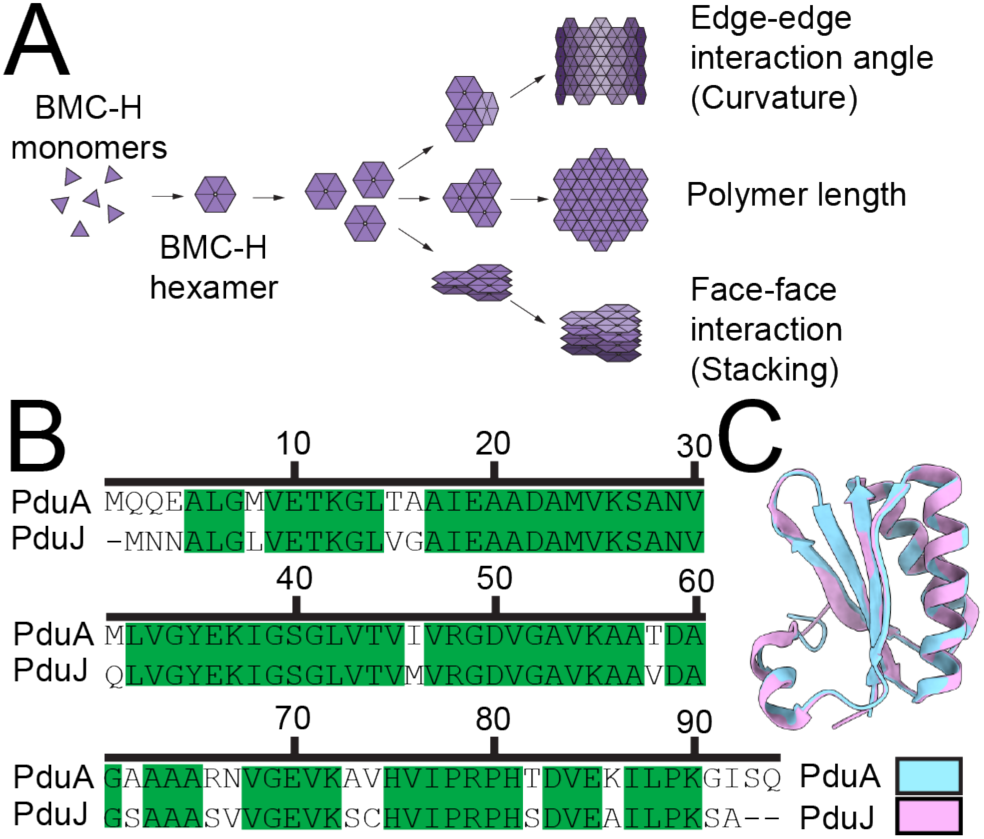
PduA and PduJ are two distinct and similar self-assembling proteins. A) Schematic flow chart describing the three types of interactions that govern BMC-H protein self-assembly. B) Sequence alignment of PduA and PduJ from *S. enterica*. Green indicates identical residues. Alignment performed using EMBOSS Needle. C) Structural overlay of PduA (light blue, PDB ID: 3NGK) and PduJ (pink, PDB ID: 5D6V). Structures were visualized using UCSF ChimeraX.

Outside of their native contexts, BMC-H proteins self-assemble to form diverse structures^12,15^. Some of the most commonly studied BMC-H proteins, including EutM from *Escherichia coli*, RmmH from *Mycobacterium smegmatis*, CsoS1A from *Halothiobacillus neapolitanus*, PduA and PduJ from *S. enterica*, and homologs from *H. ochraceum*, self-assemble into bundles of tubes when overexpressed *in vivo*^12,16–19^, in addition to nanotubes and -sheets when purified and reconstituted *in vitro*^16–20^. These self-assembled BMC-H structures have already been developed into materials with various potential applications. For example, the BMC-H protein EutM from *E. coli* was engineered into nano-scale scaffolds for enzyme co-immobilization to improve rates of *in vitro* catalysis^9^. EutM has also been engineered to form macro-scale materials upon introduction of an inducible green fluorescent protein (GFP) crosslinker^21^.

Two BMC-H proteins, PduA and PduJ, from the *S. enterica* Pdu MCP are particularly interesting candidates for engineering supramolecular protein assembly. PduA and PduJ hexamers form robust nanotube structures upon *in vivo* self-assembly. These nanotubes are so mechanically stable that they prevent cell division, leading to a linked-cell phenotype that can be upwards of 10 cells long^12,16^. When evaluated *in vitro*, PduA and PduJ spontaneously self-assemble into structures larger than the cell without the need for chemical crosslinking or genetic modifications^16,20^. PduA and PduJ’s propensity for self-assembly and the robust nature of their conferred structures renders them excellent candidates for rational design of engineered protein-based materials.

PduA and PduJ share 77.7% sequence identity (Fig. 1B) and a root mean square deviation between their structures of 0.292 Å (Fig. 1C), yet even these small differences lead to large impacts on structures formed by PduA and PduJ^22^. Prior work indicates that PduA and PduJ are functionally redundant in their ability to drive Pdu MCP assembly, which is hypothesized to be part of their native biological function^16^. However, Trettel and Winkler (2023) observed differences in PduA and PduJ self-assembly *in vitro*, in which purified PduA formed exclusively nanotubes, whereas PduJ assembled into nanotubes and hexagonal sheets. This suggests that, outside the confines of the cell, small differences in amino acid sequence may lead to differences in the morphology of the large protein assemblies. However, the requirement for purification and reconstitution of individual mutants has prevented a systematic study of the impact of sequence on supramolecular assembly.

Cell-free protein synthesis (CFPS) has the potential to overcome the limitations of cell-based protein expression by eliminating the need for labor-intensive protein purification and reconstitution. CFPS systems are capable of carrying out protein production without the need for living cells^23,24^. Instead, purified cell extract containing all required transcription and translation machinery is combined with cofactors and plasmid DNA encoding a gene of interest, allowing transcription and translation into protein fully *in vitro*^25^. CFPS is a uniquely advantageous tool for evaluating self-assembling proteins, as protein synthesis is immediately followed by assembly in the same vessel. This enables protein assembly to be evaluated as protein is synthesized, which is not feasible with other *in vitro* methods that require protein purification. Moreover, this method allows for screening tens of protein variants on the timescale of a few hours. Further, CFPS offers the additional advantage of increasing the size of the “container” from *E. coli* cells (∼0.6 fL)^26,27^ to the dimensions of the entire CFPS reaction (40-200 µL), which has implications for the assembly size as well.

Here, we demonstrate that PduA and PduJ spontaneously self-assemble into structures nearly a millimeter in length when expressed in CFPS. To do this, we incubated CFPS reactions expressing PduA-FLAG or PduJ-FLAG (herein: PduA^FL^ and PduJ^FL^) with a fluorophore-conjugated anti-FLAG antibody (Fig 2A, left). The antibody allowed us to visualize the protein assemblies on a super-resolution microscope, distinguishing PduA^FL^ and PduJ^FL^ from other cellular machinery upon fluorescent imaging. By incubating the CFPS reactions in microscope-compatible chambers, we eliminated the need to pipette or transfer the samples, which could potentially disturb the supramolecular interactions holding them together. With this method, we were able to evaluate the morphology of self-assembled PduA or PduJ structures in three dimensions (Fig. 2A). We found that PduA and PduJ rapidly and spontaneously self-assemble from single ∼9 kDa subunits into micrometer- and even millimeter-scale structures. We further demonstrate that single amino acid substitutions yield vast changes in morphology, and that other BMC-H proteins with vastly different amino acid sequences (*e.g.,* a PduA homolog from *Clostridium autoethanogenum* (*C. auto*) with 50% amino acid identity) can still self-assemble into similarly large structures. Finally, we underscore the robust nature of these proteins, demonstrating that they assemble even after lyophilization and storage at a wide range of temperatures, a key feature for their potential utility. Taken together, we show BMC-H proteins self-assemble in under an hour post-translation into materials with a wide array of morphologies and sizes, with genetic programmability.

**Figure 2.**
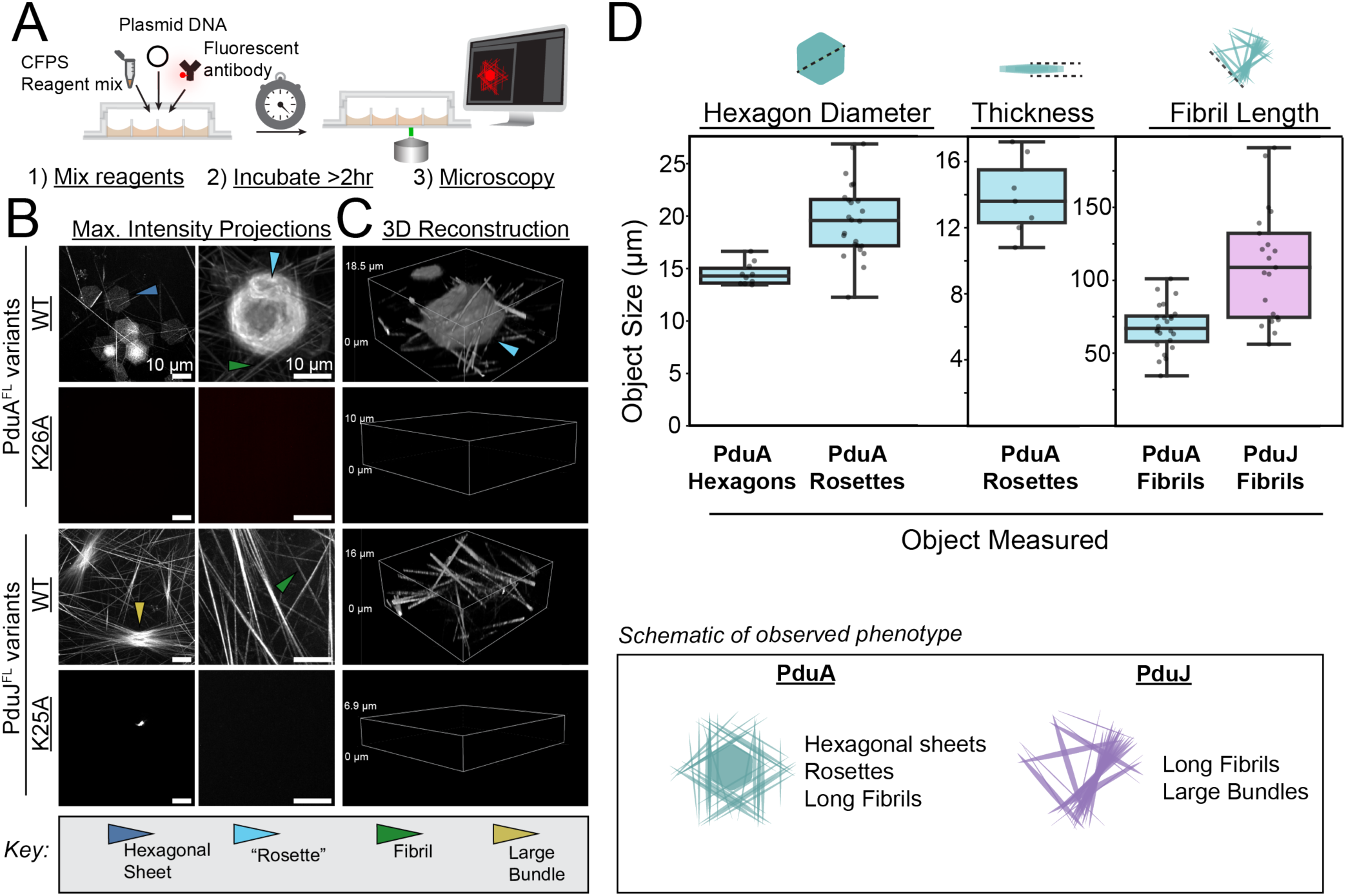
PduA and PduJ form different supramolecular structures when expressed using CFPS. A) Schematic overview of experimental design. Cell-free protein synthesis reactions containing a plasmid to express the gene of interest are co-incubated with a fluorophore-conjugated anti-FLAG antibody. Reactions are set-up in the imaging chamber and incubated for > 2 hr at 30 °C before imaging. B) Maximum intensity projections of z-stack images of CFPS reactions expressing PduA^FL^ and PduJ^FL^ variants. All scale bars = 10 µm. C) 3D reconstructions of Z-stack images using Alpha Blending. All bound boxes are cropped to 36 µm x 36 µm in XY. The heights of bound boxes in Z are labeled on the left sides of the bound boxes. D) Box and whisker plots quantifying key structural parameters of the sample, including hexagon diameter, thickness, and fibril length.

## Results/Discussion

### Cell-free–produced PduA and PduJ spontaneously self-assemble into structures on the millimeter scale

First, we aimed to determine if PduA and PduJ expressed in CFPS recapitulate previous findings that PduA and PduJ self-assemble into unique structures *in vitro.* To this end, we expressed PduA^FL^ and PduJ^FL^ in CFPS and used multiple microscopy techniques to visualize the structures. Ultimately, we found that addition of a fluorescently labeled anti-FLAG antibody into our reaction mixture yielded the clearest images of assembled structures. As described above, this setup allows specific visualization of PduA^FL^ and PduJ^FL^ assembly via super-resolution microscopy. The resulting micrographs revealed that PduA and PduJ spontaneously self-assembled into structures on the scale of tens to hundreds of microns, with the largest spanning beyond the entire field of view of the microscope (Fig. 2C-D). The structures formed in CFPS were much larger—over 0.2 mm long—than those reported by Trettel and Winkler (i.e. approximately 5 µm) who originally found that purified PduA and PduJ self-assemble into unique morphologies *in vitro*^20^.

When expressed in CFPS, PduA^FL^ formed at least four distinct micron-scale structures: 1) regular hexagonal-shaped sheets 13.5 – 16.6 microns in diameter (Fig. 2B, blue arrow, Fig. 2D); 2) a network of high-aspect-ratio structures (herein termed “fibrils”) up to 100 microns long (Fig 2B, green arrow, Fig. 2D); 3) “rosette”-like assemblies, which appeared to be stacks of hexagonal sheets upwards of 27 microns in diameter and ranging in thickness from 10 to 17 microns (Fig. 2B, light-blue arrow, Fig. 2D); and 4) protein clusters with ribbon-like assemblies (Fig. S1A, blue arrow). While fibrils appeared throughout the sample, a large fraction of these fibrils oriented parallel to and along the edges of rosettes, extending beyond the rosette perimeter and creating a structure resembling a six-pointed star with a regular hexagon in the middle.

To assess the size to which a single PduA structure could reach, we diluted PduA^FL^ 1:5 in water and used brightfield imaging at 20X magnification, which provided a large field of view. The fibrils observed under these conditions were nearly 0.6 mm long (Fig. S1B). Taken together, these data suggest that PduA^FL^ self-assembles in CFPS such that PduA hexamers interact at multiple angles, some of which allow for formation of planar hexagonal sheets while others allowed for formation of nanotubes, which collected together to form fibrils. Additionally, the large rosettes indicate that many hexagonal sheets are able to stack atop one another, suggesting hexamers not only interact through edge-edge interactions, but through face-face interactions.

Expression of PduJ^FL^ in CFPS resulted exclusively in high-aspect ratio fibril structures, some of which extended beyond the approximately 700 x 700 µm field of view on the super-resolution microscope. This contrasts with PduA^FL^ fibrils, which can be visualized end-to-end within the frame of view of the microscope. PduJ^FL^ fibrils bundled parallel to each other in many points. Interestingly, larger bundles were surrounded by fibrils that appear to splay outward from the central tightly packed point, resembling dry noodles sticking out of a bottle, where the bottleneck causes a point of tighter bundling (Fig. 2C, yellow arrow). This indicates PduJ hexamers assemble with an edge-edge interaction angle that allows fibrils to form. Overall, PduA fibrils tended to terminate around 100 microns in length, whereas most PduJ fibrils were nearly 0.2 mm long (Fig. 2D).

To confirm that edge-edge interactions are critical for PduA and PduJ assembly, we performed CFPS with non-assembling controls, PduA-K26A^FL^ and PduJ-K25A^FL^, which feature a lysine-to-alanine mutation that disrupts key electrostatic interactions that stabilize the hexamer-hexamer edge-edge interaction. These mutations completely ablate native MCP assembly *in vivo*^11^. The native Pdu MCP can tolerate a gene knockout of PduA or PduJ because they are functionally redundant, but surprisingly, replacing either PduA or PduJ with PduA-K26A^FL^ or PduJ-K25A^FL^ prevents MCP formation even when an assembly-competent counterpart is still present^11^. Upon expression of PduA-K26A^FL^ and PduJ-K25A^FL^ in CFPS, no assembled structures were detected (Fig. 2B, C), indicating that neither PduA-K26A^FL^ nor PduJ-K25A^FL^ were able to assemble into ordered structures that could be resolved by the microscope. While some aggregates were observed, similar aggregates appear in CFPS reactions prepared without adding an expression plasmid, indicating they are likely not Pdu protein assemblies (Fig. S1C). This confirms the structures we see for PduA^FL^ and PduJ^FL^ require the assembly-competent protein to be expressed in the CFPS reaction. Additionally, this indicates that edge-edge interactions are necessary for nucleation prior to face-face interactions (stacking). To confirm that the observed phenotypes for PduA-K26A^FL^ and PduJ-K25A^FL^ were not due to a lack of protein expression in CFPS, we quantified the amount of protein produced by the CFPS reactions for all constructs using C-14 leucine incorporation. This technique specifically quantifies protein produced in the cell-free reaction, as isotopically labeled leucine is only provided upon setup of CFPS. The resultant autoradiograms indicate that PduA-K26A^FL^ and PduJ-K25A^FL^ express at similar levels to PduA^FL^ and PduJ^FL^ (Fig. S1D-G). These data indicate that the lack of large protein assemblies in PduA-K26A^FL^ and PduJ-K25A^FL^ are due to the protein sequence, not their expression levels in CFPS.

Taken together, we find that PduA^FL^ and PduJ^FL^ each form unique micron-scale structures orders of magnitude larger than the bacterial cells from which they originate. We hypothesize that the slight differences between PduA and PduJ drive the observed changes in supramolecular assembly by changing both preferred hexamer interaction angles and face-face interactions between hexamers.

### Single amino acid substitutions reshape PduA assembly

Recognizing that PduA and PduJ self-assemble into structures with unique morphologies—despite being only 13 amino acids different from one another—we hypothesized that single amino acid substitutions in PduA or PduJ could also alter assembly morphology. This hypothesis is supported by work from Sinha *et al.*, which showed that specific PduA point mutants alter the morphology of Pdu MCPs^11^. Of the mutants explored in their work, we selected PduA-D59A^FL^ and PduA-K55A^FL^ to study using our CFPS system, as we expected that eliminating charge on the surface of the self-assembling subunit, PduA, would yield the greatest changes in assembly. Importantly, because residues 55 and 59 are in close proximity on the PduA surface, these two mutants also allowed us to determine if any observed differences are due to the respective residues’ charge or their position on the protein surface (Fig. 3A).

**Figure 3.**
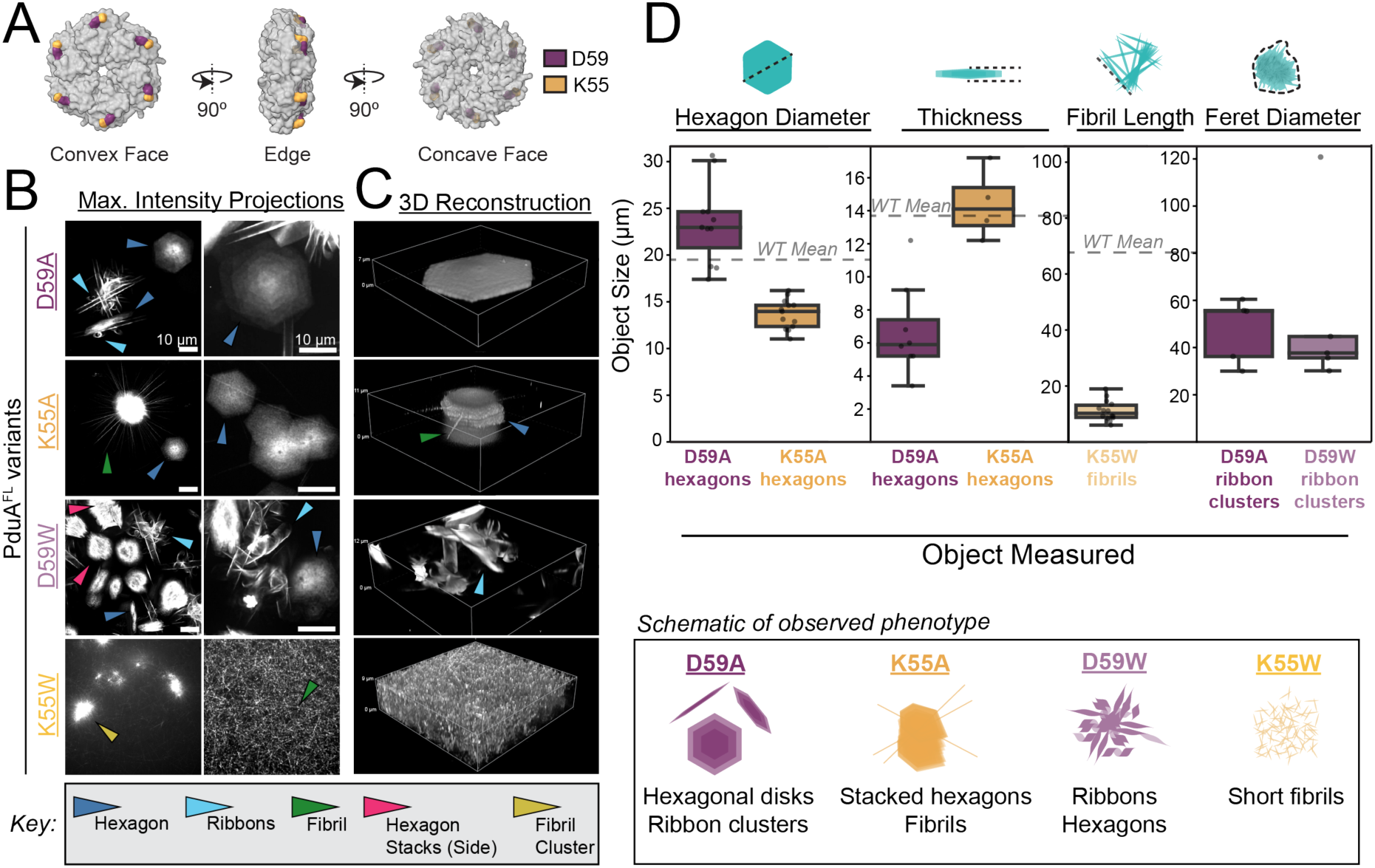
Altering charges surface residues on PduA changes supramolecular assembly. A) Structure of PduA hexamer (PDB ID: 3NGK) showing convex face (top), hexamer edge (middle), and concave face (bottom). Residue D59 is highlighted in yellow and K55 is in black. Structures were visualized using UCSF ChimeraX. B) Maximum intensity projections of z-stack images of PduA^FL^ variants imaged by confocal microscopy. All scale bars = 10 µm. C) 3D reconstructions of Z-stack images from confocal microscopy using Alpha Blending. All bound boxes are cropped to 36 µm x 36 µm in XY. The heights of bound boxes in Z are labeled on the left sides of the bound boxes. D) Box and whisker plots quantifying relevant structural parameters of the sample, including hexagon diameter, thickness, fibril length, and feret diameter of clusters.

Upon expression in CFPS, PduA-D59A^FL^ assembled into two distinct morphologies: (1) regular hexagons between 17-30 microns in diameter (Fig. 3B-C, blue arrow, Fig. 3D) and (2) clusters of protein resembling ribbons intersecting at a central point (Fig. 3B, light-blue arrow, Fig. 3D). Interestingly, in contrast to wild-type PduA^FL^, there were no “fibrils” observed in any PduA-D59A^FL^ samples. Many of the hexagons formed by PduA-D59A^FL^ appeared to have more protein in the middle and less toward the edges, as indicated by the brighter signal toward the center. We also noted that the PduA-D59A^FL^ hexagons are 6 microns thick on average (Fig. 3C) — less than half the wild-type PduA^FL^ rosette average. These data suggest that PduA-D59A^FL^ assembles through both edge-edge interactions and face-face interactions, but with weaker face-face interactions than wild type (Fig. 3D). All in all, these observations indicate that a single substitution to alanine at residue 59 can shift the micron-scale structures formed by PduA from high-aspect ratio structures to regular hexagonal disks and ribbons.

PduA-K55A^FL^ also assembled into a heterogeneous mixture of structures, fibrils and hexagons, distinct from those formed by wild-type PduA and PduA-D59A (Fig. 3B and C). with different morphologies. It formed fibrils (Fig. 3B and C, green arrow) and hexagons, contrasting with PduA-D59A^FL^. Interestingly, the fibrils formed by PduA-K55A^FL^were oriented differently than those formed by the wild-type PduA^FL^. While wild-type PduA^FL^ self-assembled into fibrils that aligned tangentially to the rosettes, PduA-K55A^FL^ fibrils intersected in the center of the hexagonal rosette (Fig. 3B and C, green arrow), appearing as though they passed through the center of the thicker stack of hexagons. The PduA-K55A^FL^ fibrils also only appeared in proximity to the rosettes and did not form a matrix throughout the entire sample, as the WT PduA^FL^ fibrils did. Unique to PduA-K55A^FL^, some hexagonal stacks were taller in Z than their diameters in XY; these stacks were up to 17 microns in height, and the individual hexagonal sheets comprising them (Fig. 3C, blue arrow) were under 16 microns in diameter (Fig 3D). This suggests that mutating the charged lysine at position 55 to alanine likely increases the face-face interactions between PduA hexamers, promoting sheets to stack higher compared to the wild type. These findings also indicate the charge and size of lysine 55 and aspartic acid 59 are significant for mediating supramolecular PduA self-assembly.

By changing two charged residues to hydrophobic alanine residues, we were able to shift PduA self-assembly to new morphologies. Next, we sought to disrupt these assemblies entirely by mutating these two residues to tryptophan, the largest amino acid. We hypothesized that mutation to tryptophan would cause steric hindrance, blocking stacking interactions and preventing larger-scale assembly. To test this hypothesis, we cloned two new variants, PduA-D59W^FL^ and PduA-K55W^FL^. Imaging of CFPS reactions expressing these mutants revealed that they assemble differently both from the wild type and from the alanine mutants. Neither PduA-D59W^FL^ nor PduA-K55W^FL^ completely ablate PduA assembly like the K26A mutant; rather, these mutations drive formation of previously unreported morphologies.

PduA-D59W^FL^ assembled into a heterogenous mixture of clustered, loosely packed hexagons (Fig. 3B and C, blue and pink arrows) and clusters of curled ribbons (Fig. 3B and C, light-blue arrows). The PduA-D59W^FL^ hexagons were not as tightly and regularly packed as the PduA-D59A^FL^ hexagons, leading to what looked like haystack when viewed from the side (Fig 3B, pink arrows). Additionally, PduA-D59W^FL^ assembled into clusters of ribbons similarly to PduA-D59A^FL^ (Fig. 3B and C, light-blue arrow). The largest clusters in PduA-D59W^FL^ reached sizes over 120 µm in diameter, over twice as large as any clusters observed for PduA-D59A^FL^ (Fig. 3D, S3A). These did not appear to be random aggregates as they had similar features (mostly aggregates of ribbons) each time they appeared. The ribbon clusters were consistent in appearance in each replicate on different days, and they did not appear in any negative controls (PduA-K26A^FL^ or PduJ-K25A^FL^ (Fig. 2B and C)).

Mutating lysine at residue 55 to tryptophan (PduA-K55W^FL^) caused a radical shift from all previously observed assemblies. PduA-K55W^FL^ predominantly assembles into short fibrils between 6 to 18 microns in length (Fig. 3D), though some clusters appeared throughout the sample (Fig. 3B, yellow arrow). We hypothesize that the shift to shorter fibrils is a result of decreased edge-edge interaction strength, while the fibril shape is dictated by the angle of edge-edge interactions between PduA hexamers.

Overall, these results demonstrate that single amino acid substitutions can completely redirect supramolecular PduA self-assembly, limiting the length or altering the organization/orientation of assembled structures. Additionally, these data demonstrate that residues not directly involved with the interaction at the hexamer-hexamer interface (canonically, K26, N29, R79)^11^ are essential for regulating the orientation and strength of interactions between PduA hexamers.

### Pore residues play a critical role in PduA self-assembly in CFPS

Since mutations on the PduA surface and on the hexamer edge caused noticeable changes in supramolecular assembly and structure morphology, we hypothesized that mutations to other regions in PduA could also impact self-assembly. To this end, we investigated mutations near the PduA hexamer pore, which is demonstrably mutable *in vivo*—mutations to residues in the pore do not disrupt native Pdu MCP assembly^28–31^. The pore thus presents an interesting target for interrogating how small sequence changes that do not impact assembly in the native system impact large-scale PduA assembly. We evaluated S40 and L42 because of their proximity to the pore (Fig. 4A), and tested alanine and tryptophan mutants (S40A, L42A, S40W, and L42W) for impact on PduA assembly due to the differences in the sizes of these residues and how they can drive assembly.

**Figure 4.**
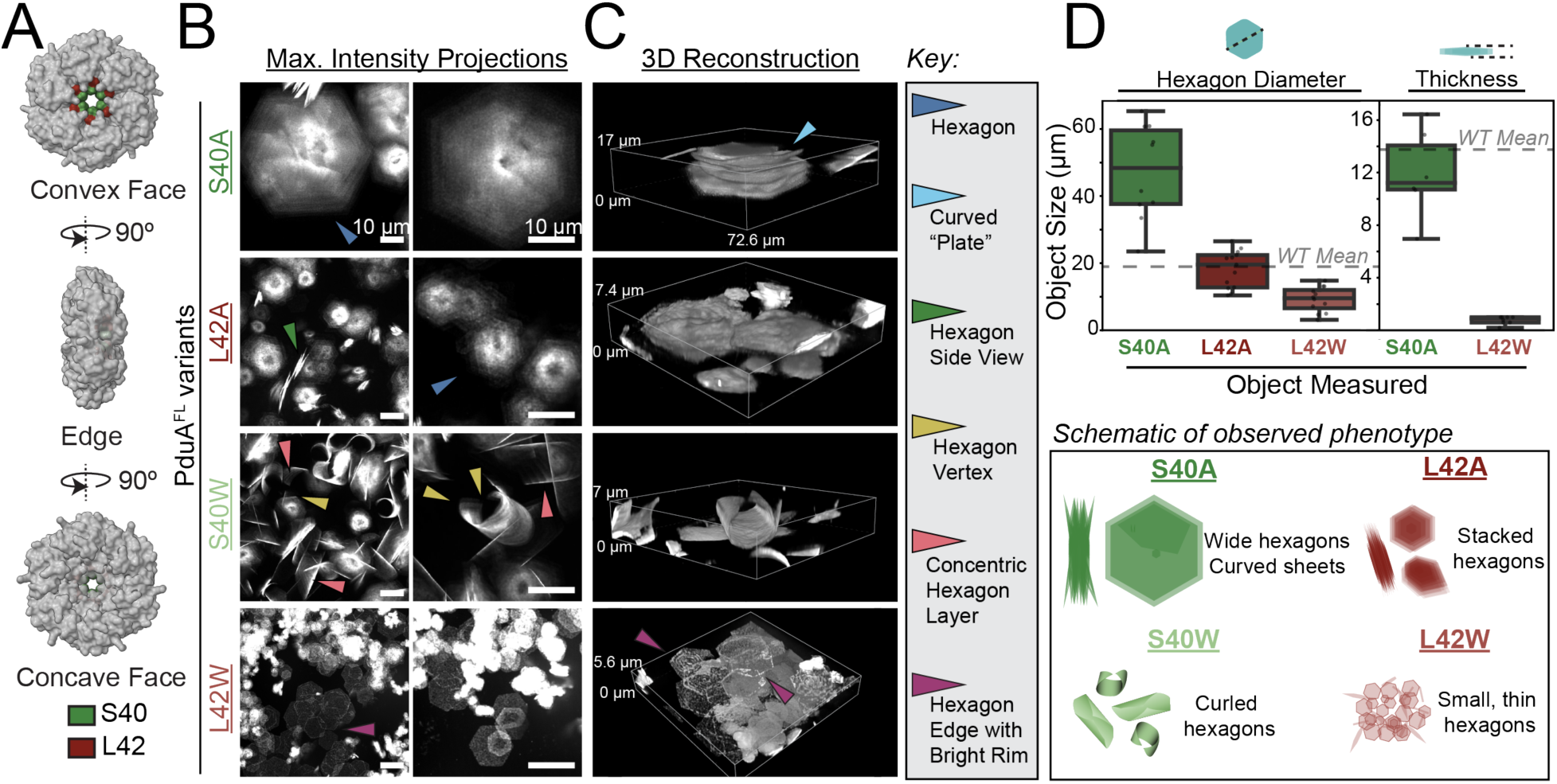
Altering PduA pore residues changes supramolecular assembly. A) Structure of PduA hexamer (PDB ID: 3NGK) showing convex face (left), hexamer-hexamer interface (middle), and concave face (right). Residue S40 is highlighted in green and L42 is in brown-red. Structures were visualized using UCSF ChimeraX. B) Maximum intensity projections of z-stack images of PduA^FL^. All scale bars = 10 µm. C) 3D reconstructions of Z-stack images using Alpha Blending. The bound box for S40A (top) is cropped to 72.6 µm x 72.6 µm in XY. All others are cropped to 36 µm x 36 µm in XY. The heights of bound boxes in Z are labeled on the images, left of the bound boxes. D) Box and whisker plots quantifying relevant structural parameters of the sample, including hexagon diameter and hexagon thickness.

We first tested S40 as it is the furthest residue from the hexamer-hexamer interface. A mutation to alanine in this position has previously been demonstrated to impact diffusion of substrates through the native MCP shell but has not been characterized in respect to assembly^28,31^. Mutation of S40 to alanine in PduA once again shifted the distribution of observed structures in CFPS—PduA-S40A^FL^ forms no fibrils and instead forms only regular hexagon-shaped materials (Fig. 4C, blue arrow). Further, hexagons formed by PduA-S40A^FL^ differ from those formed by PduA-D59A^FL^ and PduA-D59W^FL^ in that PduA-S40A^FL^ hexagons had greater diameters—the smallest was 25 microns from one vertex to another, which is over 6 microns larger than the wild-type mean. Moreover, the largest we observed were 65 microns in diameter (Fig. 4B-D), over 3 times the diameter of the wild-type average. Viewing these hexagons from the side (Fig. 4C) highlighted that the structures are formed from many hexagonal sheets that splay out from a central, densely packed point. The structures were between 7-16 microns thick (Fig. 4D), and the individual layers seemed to be curved like a shallow plate (Fig 4C, light-blue arrow). The center point of the S40A hexagons appeared darker on the micrographs, which we believe occurred because to the protein was packed too tightly for the antibody stain to access. These results demonstrate that PduA hexamer pore residues are able to modulate assembly.

To see if other pore residues impact assembly, we evaluated L42, as it is the nearest residue to S40 on the convex face of the hexamer (Fig. 4A). PduA-L42A^FL^ also spontaneously assembled into hexagons, and we did not observe any fibrils within the resolution limit of the microscope (Fig. 4B). The edges of these hexagons were not as even as those of the PduA-S40A^FL^, PduA-K55A^FL^, or PduA-D59A^FL^ mutants, similar to the PduA-D59W^FL^ hexagons in the maximum intensity projections. PduA-L42A^FL^, however, did not form any clusters of ribbons, unlike PduA-D59W^FL^. Side views of the PduA-L42A^FL^ structures (Fig. 4C, green arrow) indicated that they were not as thick as other hexagons from other PduA^FL^ variants. This demonstrates that the capacity of pore residues for regulating PduA assembly is not limited specifically to residue S40.

Since previous tryptophan mutations significantly impact PduA assembly, we decided to investigate assembly of PduA-S40W^FL^ and -L42W^FL^. We hypothesized that introduction of a bulky residue like tryptophan at the innermost point of PduA could change the curvature of the PduA hexamer, leading to a difference in assembly compared to the WT or the alanine mutants. While PduA-S40A^FL^ assembled into large, mostly planar (but slightly curved) hexagons up to 65 microns in diameter that form stacks 8-16 microns thick (Fig. 4D), PduA-S40W^FL^ formed slightly curled hexagonal sheets (Fig. 4B). When viewing PduA-S40W^FL^ through single slices, the morphology of these assemblies was not easily discernible, but maximum intensity projections and 3D reconstructions revealed shapes similar to curly, hollow corn chips or cannoli. They appeared as if two opposite vertices (Fig. 4C and D, yellow arrow) of a regular hexagon-shaped surface were folded together to make a hollow cylinder. These materials also appeared to be stacks of multiple layers of hexagonal sheets as there are rings of concentric hexagons (Fig. 4C and D, magenta arrow) with greater fluorescence intensity toward the middle of the structure and dimmer signals near the edges, similar to PduA-D59A^FL^.

In contrast, PduA-L42W^FL^ assembled into regular hexagons that were planar and much thinner than any other variant visualized thus far. While all other regular hexagons we observed demonstrated some degree of perceived stacking interactions, we did not observe that with PduA-L42W^FL^. These hexagons were thin with a bright rim around the edges (Fig 4B and C, purple arrow). Most of these hexagons were barely perceptible within one Z slice on the microscope and required a Z- or 3D-projection to resolve the structures.

All eight variants characterized in Figures 3 and 4 formed different supramolecular structures from wild-type PduA^FL^, whether it was the disappearance of fibrils, the thickness of hexagonal shaped rosettes, or the curvature of the protein sheets that formed. This underscores that these four residues play a critical role in the assembly of PduA structures. Additionally, these results indicate that residues across the PduA structure, whether they are on the hexamer edge, face, or near the pore, can regulate PduA assembly and the resultant morphology and size of structures in ways that are not obvious *a priori*.

### Mutations to PduJ underscore the ability to engineer molecular assembly and highlight differences with PduA

Since the four PduA residues we tested in previous sections are conserved in PduJ, we decided to analyze their impacts on PduJ assembly. We hypothesized that these mutations would yield similar morphology changes in PduJ as they did in PduA, as PduA and PduJ have nearly identical primary, secondary, and tertiary structures. To test this hypothesis, we investigated eight PduJ^FL^ variants: PduJ-D58A^FL^, -K54A^FL^, -D58W^FL^, - K54W^FL^, -S39A^FL^, -L41A^FL^, -S39W^FL^, and -L41W^FL^. Interestingly, out of eight mutants, only one (PduA-K55W^FL^/PduJ-K54W^FL^) assembled identically between PduA and PduJ (Fig. 5A and B). The other mutants underscored the difference in assembly properties between PduJ and PduA, as the 13 differing residues between PduA and PduJ continue to drive assembly toward different morphologies.

**Figure 5.**
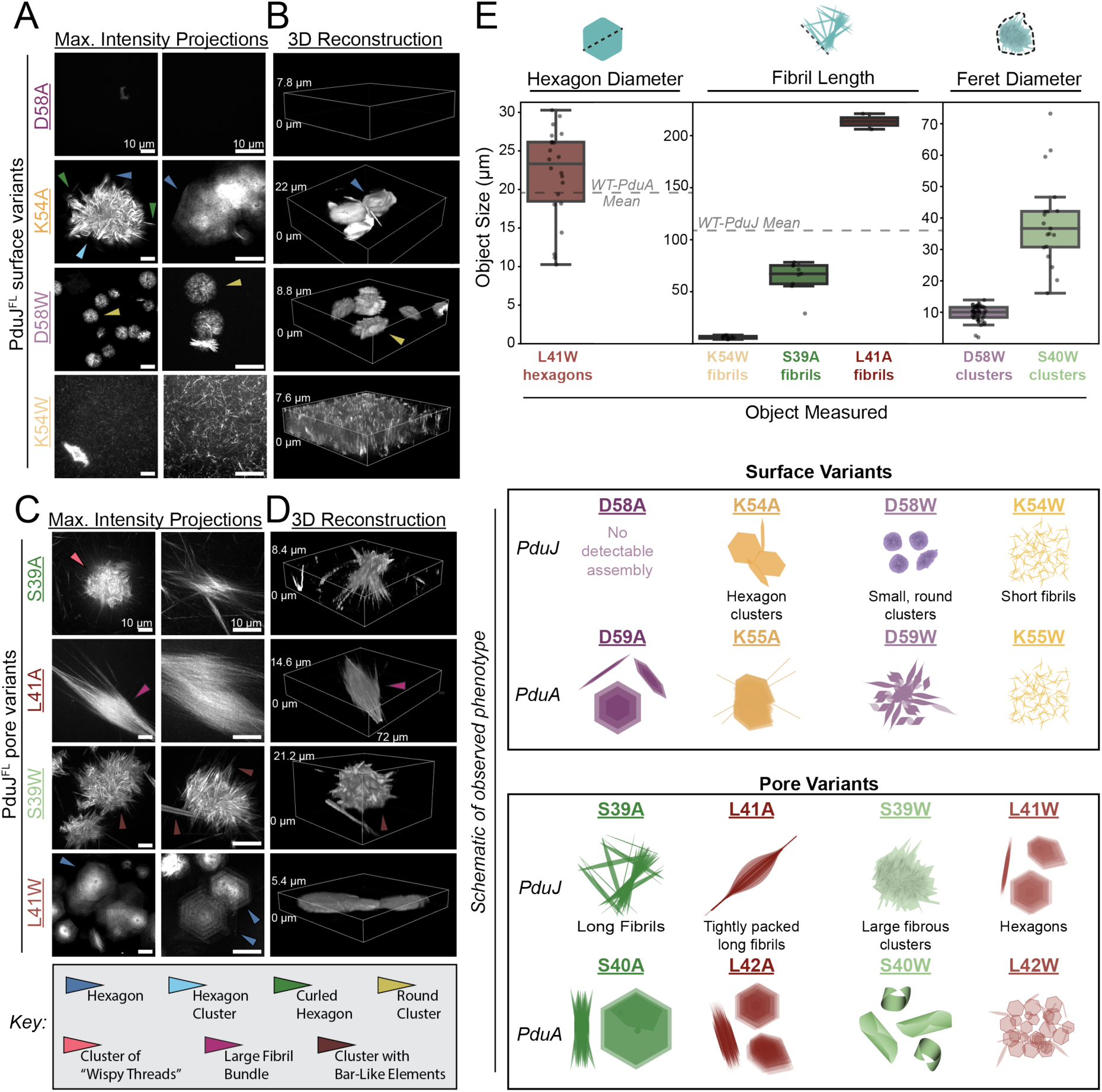
Mutations to PduJ reveal key residues that govern both PduA and PduJ supramolecular assembly. A) Maximum intensity projections of z-stack images of PduJ^FL^ surface variants. All scale bars = 10 µm. B) 3D reconstructions of Z-stack images using Alpha Blending. All images are cropped to 36 µm x 36 µm in XY. The heights of bound boxes in Z are labeled on all images to the left of the bound boxes. C) Maximum intensity projections of z-stack images of PduJ^FL^ pore variants. All scale bars = 10 µm. D) 3D reconstructions of Z-stack images using Alpha Blending. The bound boxes are cropped to 36 µm x 36 µm in XY, except the bound box for L41A (third from bottom) is cropped to 72 µm x 72 µm in XY. The heights of bound boxes in Z are labeled on all images to the left of the bound boxes. E) Box and whisker plots quantifying relevant structural parameters of the sample, including hexagon diameter, fibril length, and ferret diameter of clusters.

PduJ-D58A^FL^ did not assemble into structures perceptible within the resolution limit of the microscope (XY: 120 nm) (Fig. 5A and B). This contrasted greatly with PduA-D59A^FL^, which formed regular hexagonal disks and clusters of ribbons or helices. PduJ-K54A^FL^ formed hexagons (Fig. 5A and B, blue arrow) that were similar to the free-floating hexagons in PduA-K55A^FL^, but the PduJ variant did not form any fibrils. PduJ-K54A^FL^ was also not as homogenous as PduA-K55A^FL^; PduJ-K54A^FL^ hexagons sometimes clustered together into larger aggregates (Fig. 5A, light-blue arrow) and occasionally the hexagons were slightly curled (Fig. 5A, S3A, green arrow), something we did not see in its PduA counterpart.

Knowing that PduJ-D58A^FL^ did not assemble into any structures we could see on the microscope, we hypothesized that PduJ-D58W^FL^ would also not form visible structures, since alanine and tryptophan both contain hydrophobic side chains. Instead, PduJ-D58W^FL^ assembled into round clusters (Fig. 5A and B, yellow arrow) with a morphology not observed in any PduA variants. These clusters did not appear hexagonal but were consistent in appearance each time they occurred. They were between 4-7 microns thick and 6-14 microns in diameter (Fig. 5E) and appeared to have an uneven surface, with brighter and dimmer signals across the cluster surface. Next, we imaged PduJ-K54W^FL^, which had the same appearance as PduA-K55W^FL^ (Fig. 5B). This was a surprising result as we had not yet seen any two variants give rise to the same morphology in our study. These data indicate that the charged lysine in PduJ position 54 (PduA position 55) is critical for regulating PduA and PduJ self-assembly. A mutation to tryptophan in this position likely forces a bent edge-edge interaction angle, leading to a fibril phenotype, while also weakening the strength of edge-edge interactions, leading to shorter fibril length (5 microns on average) compared to wild type (110 microns on average) (Fig 5D).

Because the four surface mutants we evaluated in PduA (D59A, K55A, D59W, and K55W) led to morphology changes in PduA assemblies, we hypothesized that mutations to pore residues would alter PduJ assembly as well. To that end, we expressed and imaged PduJ-S39A^FL^, the PduJ mutant analogous to PduA-S40A^FL^. Since this mutation caused PduA to self-assemble into hexagonal sheets that were 65 microns in diameter and stacked over 10 microns tall (Fig. 4D), we hypothesized PduJ-S39A^FL^ would demonstrate an increased propensity for hexamer stacking and elongation. Instead, PduJ-S39A^FL^ formed structures similar to wild-type PduJ^FL^. This mutant almost exclusively formed long fibrils between 30-80 microns long (compared to wild type 56-190 microns) (Fig. 5E) and bundling as wild-type PduJ^FL^ did (Fig. 2B-C). There were, however, some instances in which PduJ-S39A^FL^ assembled into clusters of wispy threads (Fig 5C, pink arrow).

Because all mutations to PduA position 42 studied led to hexagonal shaped assemblies, we hypothesized that the PduJ-L41 mutants would similarly assemble into hexagonal structures. PduJ-L41A^FL^, however, assembled into long fibrils over 200 microns in length that extend beyond the field of view of the microscope. The positions of wild-type PduJ^FL^ fibrils appeared mostly stochastic throughout the sample, whereas PduJ-L41A^FL^ fibrils usually appeared together as a very tight bundle of thin hair-like structures (Fig. 5C and D, purple arrow). The diameter of these bundles ranged between 11-20 microns across and either tapered off away from the center of the bundle and/or splayed outward away from the nucleus of the bundle (Fig. 5C-D, Fig. S3B).

Next, we evaluated PduJ-S39W^FL^. In PduA, this mutation led to formation of curled hexagonal sheets, resembling cannoli shells. In contrast, PduJ-S39W^FL^ formed into spherical clusters with many bar-shaped protrusions (Fig. 5C and D, brown arrow) pointing outward from a central point (Fig 5D and E). The clusters were between 25 microns to 50 microns across and resembled materials made by Zhang et al. (2024) out of EutM, a different BMC-H protein^21^.

Finally, we analyzed PduJ-L41W^FL^. PduJ-L41W^FL^ self-assembled exclusively into hexagonal sheets between 10 and 30 microns in diameter. Thinner layers of the sheets appeared to be stacked on top of one another in the maximum intensity projections as there appeared to be concentric hexagons, with a brighter signal approaching a dark center point and dimmer signal from the concentric hexagonal rings away from the center (Fig. 5C, dark blue arrows). The darker center point was likely an artifact of our imaging method, as the antibody cannot access protein if it is too tightly packed.

Overall, these data indicate that these four conserved residues (D59, K55, S40, and L42 in PduA) can have large impacts on protein assembly in both PduA and PduJ. Across the eight mutations evaluated in both PduA and PduJ, none exactly recapitulated the morphology of the wild-type assemblies. PduJ-S39A^FL^ was the most similar to PduJ^FL^ but contained clusters that we have not seen in wild-type PduJ^FL^. Additionally, only one mutation—PduA-K55W^FL^ and PduJ-K54W^FL^—gave rise to assemblies with the same morphologies in both PduA and PduJ. The observation that analogous mutations to PduA and PduJ assemble into different morphologies suggests that the 13 amino acids that differ between PduA and PduJ drive the observed differences in PduA and PduJ assembly.

### A homolog of PduA from *Clostridium autoethanogenum* spontaneously forms large, assembled structures in CFPS

Since *S. enterica* PduA and PduJ demonstrate a high propensity for self-assembly in CFPS, we hypothesized that BMC-H proteins from other MCP operons and other organisms would as well. Therefore, we expressed 4 FLAG-tagged BMC-H proteins in CFPS: (1) EutM^FL^ from *E. coli* (uniprot: P0ABF4), (2) RmmH^FL^ from *M. smegmatis* (NCBI: YP_884687), (3) CsoS1A^FL^ from *H. neapolitanus* (uniprot: P45689), and (4) CAETHG3289^FL^, a homolog of PduA from *C. auto* (uniprot: UPI0003BAD469)^32^. These BMC-H proteins range from 50.5% to 71.6% in sequence identity to PduA (Fig. S5A), and thus we expected them to self-assemble in CFPS a similar manner to PduA and PduJ. Upon imaging using the same conditions as for PduA and PduJ, we were unable to resolve any structures for EutM^FL^, RmmH^FL^, and CsoS1A^FL^ (Fig S5B). However, we did observe structures of CAETHG3289, which shares 71.6% sequence identity with PduA— the highest in similarity of the four BMC-H proteins tested (Fig 6A)^33,34^. Additionally, the AlphaFold3^35^ predicted structure of CAETHG3289 overlays closely with the *S. enterica* PduA structure and has an RMSD of 0.366 Å (Fig 6B).

**Figure 6.**
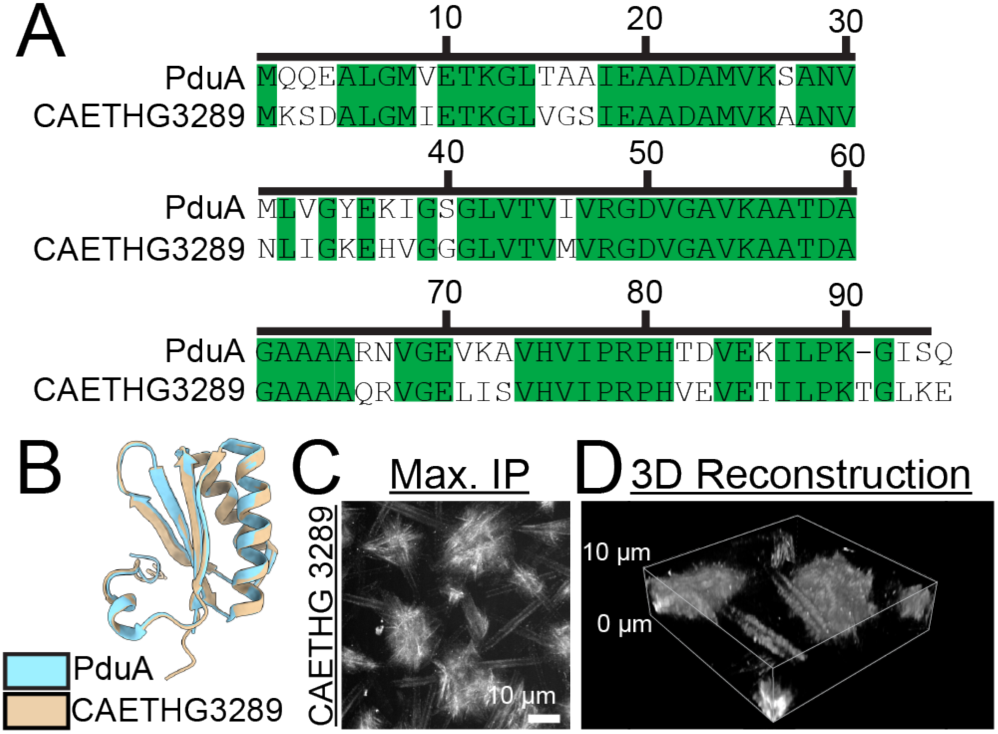
PduA homolog with only 50% similarity spontaneously self-assembles into a tens of microns-scale material. A) Sequence alignment of PduA from *S. enterica* and CAETHG3289 from *C. auto*. Green indicates identical residues. Alignment performed using EMBOSS Needle. B) Structural overlay of PduA (light blue, PDB ID: 3NGK) and CAETHG3289 (tan, alphafold). Structures were visualized using UCSF ChimeraX. C) Maximum intensity projections of z-stack images of CFPS reactions expressing CAETHG3289FL. Scale bar = 10 μm. E) 3D reconstruction of Z-stack image using Alpha Blending. Bound box is cropped to 36 μm x 36 μm in XY. The height of the bound box in Z is labeled on the image to the left of the bound box.

CAETHG3289^FL^ assembled into a mix of two morphologies. The majority of the assemblies in the samples appeared as clusters between 9 and 28 microns in diameter comprising wispy threads (Fig 6C, S5C). This scale is comparable to what we saw in many *S. enterica* PduA and PduJ variants that also lacked clearly defined fibril-like structures (e.g., the PduJ-L41W hexagon diameter ranged 10-30 microns) (Fig. S5C). In addition to these clusters, there were some higher aspect ratio structures as well, which appeared as bar-like shapes. It is unclear if these “bars” were bundles of microtubes, hollow tubes, a sheet curled in on itself like a Swiss roll, or another structure entirely. When we viewed a single slice of the image (Fig S6D), the signal appeared as two parallel lines. PduA-S40W^FL^ also had this appearance in single z-slices, however maximum intensity projections and 3D renderings revealed the hexagonal “cannoli” shapes. By contrast, the maximum intensity projections and 3D renderings of CAETHG3289^FL^ did not reveal any such structures, suggesting that these “bars” are either standalone or connected by a structure that is smaller than the resolution limit of the microscope used here, which is 120 nm (Fig 6D).

These data support that BMC-H proteins from other organisms have the potential to form large protein assemblies similar to PduA and PduJ. Although only 71.6% of residues are identical between *S. enterica* PduA and CAETHG3289, both are able to spontaneously self-assemble into a material on the scale of tens of microns in size (Fig. S5C). This result can be used to inform future studies that wish to evaluate the exact residues that are essential for self-assembly and fine-tune the supramolecular interactions that hold BMC-H structures together.

### PduA and PduJ self-assembly are robust across a wide range of temperatures and lyophilization

The PduA and PduJ structures we observe could have utility in a materials context, for example as scaffolds for enzymes, for stabilizing hydrogels, or for metal templating^9,36,37^. Doing so requires the structures maintain a stability to various post-synthesis processing steps, particularly to enable transit between manufacturing locations. To that end, freeze-drying is a common method used to decrease the mass of materials during shipment in addition to increasing shelf-life of biologics and biological tools like CFPS by minimizing hydrolysis during storage^38,39^. Therefore, we aimed to determine if our PduA and PduJ assemblies withstand the freeze-drying process. To determine the stability of PduA and PduJ to this process, we first incubated CFPS reactions expressing PduA^FL^, PduJ^FL^, PduA-K26A^FL^, and PduJ-K25A^FL^ as we did in the earlier sections but without a fluorescent antibody (Fig. 7A). After letting the reactions incubate for more than 2 hours, we freeze-dried the samples. We then placed them in storage at either 4 °C (standard refrigeration), 24 °C (standard room temperature), or 55 °C (heat) for one week. After one week, we rehydrated the samples in an antibody solution for visualization on the microscope.

**Figure 7.**
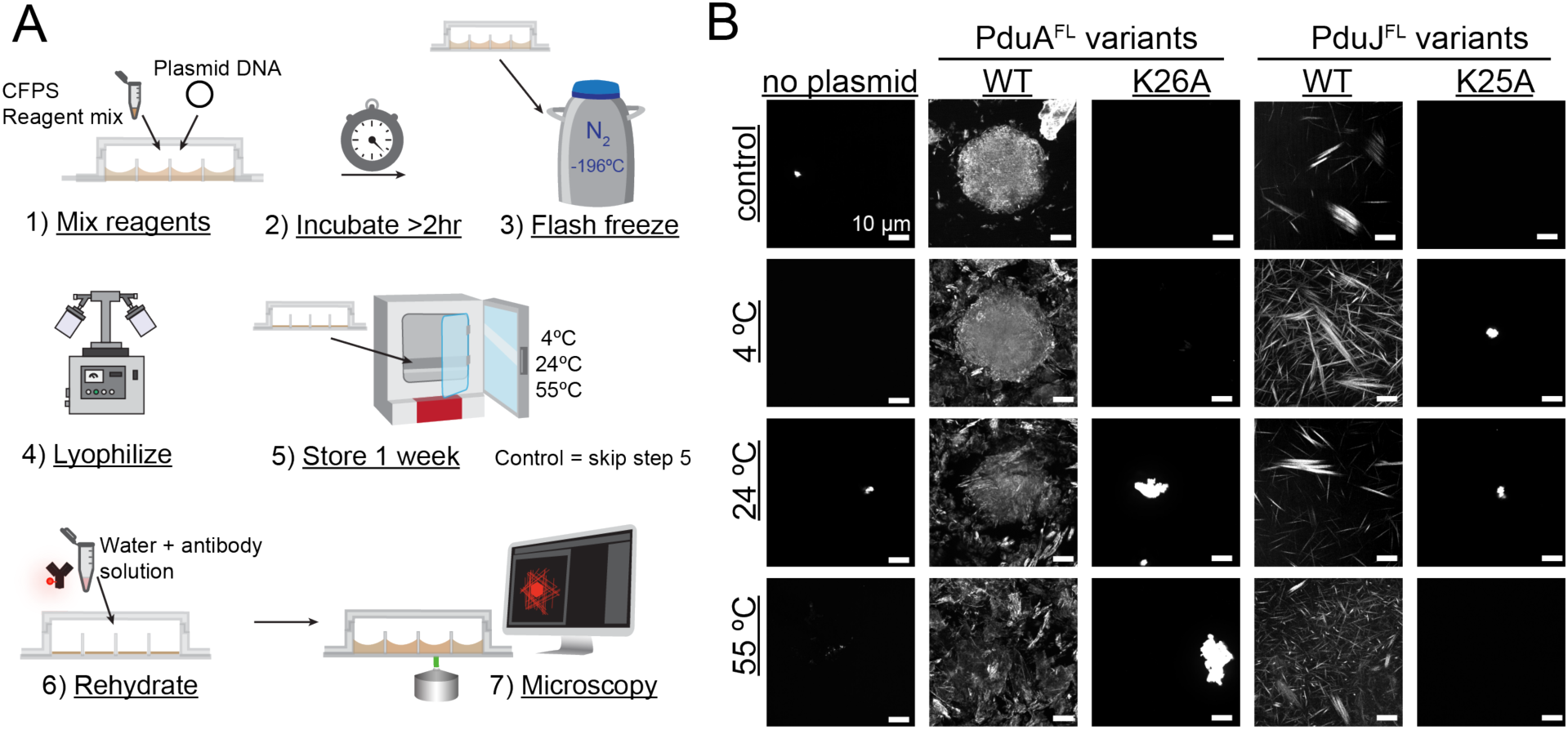
PduA and PduJ assemblies reform structures after lyophilization, heating, and rehydration. A) Schematic overview of experimental design. Cell-free protein synthesis reactions containing a plasmid to express the gene of interest are incubated for > 2 hr at 30 °C before flash freezing and lyophilization. Samples were then incubated at 4, 24, or 55 °C for one week before being rehydrated in a solution containing a fluorophore-conjugated anti-FLAG antibody for imaging. B) Maximum intensity projections of z-stack images of CFPS reactions expressing PduAFL and PduJFL variants. No plasmid indicates that no plasmid was added into CFPS reaction, producing no protein. Labels on left describe the temperature at which the sample was incubated for 1 week prior to rehydration and visualization; control was immediately rehydrated and imaged, not incubated for one week. All scale bars = 10 μm.

Here, we saw that PduA^FL^ structures remained visible after freeze-drying; however, their morphology no longer resembled the combination of fibrils and hexagonal rosettes after rehydration. They appeared as a faint hexagonal shape on the order of 40 microns (Fig. S6A) but not as distinct as they were pre-freeze drying (e.g. those in Fig. 2). To further evaluate stability over time, we stored freeze-dried PduA^FL^ samples at 4 °C, 24 °C, and 55 °C for one week. Structures continued to appear in all PduA^FL^ samples regardless of the temperature they were stored in. However, the hexagonal shapes became decreasingly visible at higher temperatures, with none persisting after 1 week at 55 °C. To confirm the structures we saw were created by PduA^FL^, we again compared the structures to our non-assembling control, PduA-K26A^FL^, and a CFPS reaction with no plasmid. PduA^FL^ structures were not present in either the “no plasmid” CFPS reaction nor the PduA-K26A^FL^ mutant.

Next, we evaluated the stability of PduJ^FL^ using the same method as previously described. Post-lyophilization, PduJ^FL^ structures were nearly identical in appearance as they were pre-freeze drying (e.g. those in Fig. 2). The structures visualized were exclusively high-aspect ratio fibrils (Fig. 7B). Since PduJ^FL^ morphology persisted upon freeze-drying and rehydration, we next incubated the lyophilized PduJ^FL^ samples at 4°C, 24°C, and 55°C for one week as we did with PduA^FL^. After one week at all of the aforementioned temperatures, PduJ^FL^ still formed high-aspect ratio structures, though they were not as long as the non-lyophilized PduJ^FL^ fibrils (Fig. S6B). Once more, comparing PduJ^FL^ fibrils to our non-assembling control, PduJ-K25A^FL^, we could confirm that the structures seen in the PduJ^FL^ samples were due to the presence of PduJ^FL^ rather than random aggregates.

Overall, PduA and PduJ not only spontaneously self-assemble into structures tens to hundreds of microns in size within 2 hours post-translation, but they are also incredibly robust materials, withstanding temperatures up to 55 °C for over one week. While PduA^FL^ morphology is impacted by the lyophilization process, PduJ^FL^ maintains its morphology post lyophilization. This demonstrates that PduA and PduJ structures remain robust post-freeze drying and incubation at elevated temperatures.

### Conclusions

In this study, we demonstrate that PduA and PduJ both self-assemble into materials hundreds of microns in size within two hours of their translation. We further demonstrate that single amino acids substitutions to PduA and PduJ sequences modulate the PduA and PduJ self-assembly process to yield millimeter-scale changes in morphology. By making single amino acid mutations in PduA and PduJ, we form materials with a diverse array of structures (Fig. 8). We hypothesize that these altered morphologies are a result of changes in the strength of the three main hexamer-hexamer interactions (edge-edge interaction strength, edge-edge-interaction angle, and face-face interaction strength), mediated by these point mutations. In addition, we show that a PduA homolog from *C. auto* with 71.6% amino acid identity to *S. enterica* PduA assembles into materials tens of microns in length, indicating that formation of large, self-assembled structures is a more general property of BMC-H proteins.

**Figure 8.**
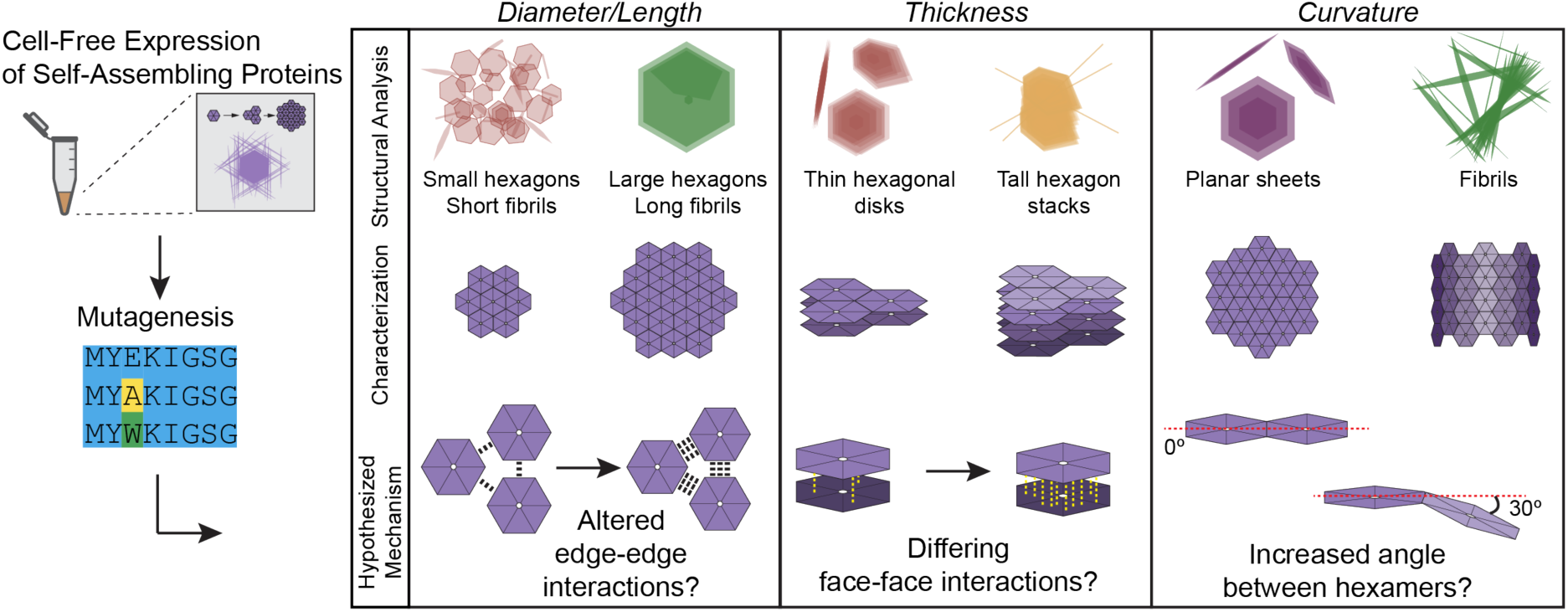
Extrapolating nano-scale interactions from micro-scale structures. Schematic overview of how we hypothesize mutations to PduA or PduJ may impact the hexamer-hexamer interactions in three ways: edge-edge interaction strength, face-face interaction strength, and edge-edge interaction angle.

We additionally show that structures assembled from PduA and PduJ deform but maintain visible supramolecular assemblies upon freeze-drying and heat exposure. This work constitutes a significant step toward the use of self-assembling proteins for biomaterial applications. We expect that future studies of these proteins and their variants will elucidate design rules that detail how different PduA and PduJ residues regulate self-assembly by modulating the three apparent hexamer-hexamer interactions.

## Acknowledgements

We gratefully acknowledge funding from the National Science Foundation (NSF) Materials Research Science and Engineering Center (MRSEC) at Northwestern University under award number DMR-2308691. This work was supported in part by the Army Research Office (W911NF-19-1-0298) and by the CBC (W52P1J-21-9-3023). Microscopy was performed at the Biological Imaging Facility at Northwestern University (RRID:SCR_017767), graciously supported by the Chemistry for Life Processes Institute, the NU Office for Research, the Department of Molecular Biosciences.

We thank all members of the Tullman-Ercek lab for their helpful discussions and support. We thank Dr. Tong Zhang from BIF at Northwestern University for his assistance with operating the Nikon SoRa.

## Methods

### Plasmids

(Table S1). All genes of interest were cloned into the pJL1 plasmid^40^ [pJL1 was a gift from Michael Jewett (Addgene plasmid # 69496; http://n2t.net/addgene:69496; RRID:Addgene_69496)]. Gene sequences were either ordered through Twist Biosciences as pre-cloned vectors, through golden gate cloning^41^, or by altering the wild-type sequences using the KOD Hot Start DNA Polymerase (Millipore Sigma Cat. No. 71086-3) for QuikChange. Oligonucleotides used for Golden Gate cloning or QuikChange can be found in Table S2. Plasmids were expressed in *E. coli* DH10B then purified using ZymoPURE II Plasmid Midiprep Kit (Zymo Research Cat. No. D4200).

### Cell Extract Preparation [A]

Unless otherwise noted, cell extracts were produced using the following method. *E. coli* BL21 (DE3) (Life Technologies) (Table S3) cells were prepared into cell-free extracts via growth, harvest, lysis, and preparation as previously described^25,42^. Briefly, cells were cultured in full-baffled flasks with 1 L of 2x YTPG (16 g/L tryptone, 10 g/L yeast extract, 5 g/L NaCl, 7 g/L potassium phosphate monobasic, 3 g/L potassium phosphate dibasic, 18 g/L glucose) at 37 °C and shaking 225 RPM. T7 RNA Polymerase production in the cultures was induced using 1 mM isopropyl-β-D-thiogalactopyranoside (IPTG) when cultures reached an OD_600_ of 0.6-0.8. Then, at an OD_600_ of 3.0 ± 0.1, cells were harvested via centrifugation at 5,000 x *g* for 10 minutes at 4 °C. The supernatant was removed from the cells and the remaining pellets were washed three times with cold S30 buffer (10 mM tris acetate, pH 8.2, 14 mM magnesium acetate, and 60 mM potassium acetate). Each wash was followed by centrifugation at 10,000 x *g* at 4 °C. After the last wash, cells were flash frozen with liquid nitrogen and stored at - 80 °C. In preparation for lysis, frozen cell pellets were thawed on ice for 1 h before resuspension in cold S30 buffer. Cells were then lysed using the EmulsiFlex-B15 homogenizer (Avestin) in a single pass at a pressure of 20,000-25,000 psi. Remaining cell debris was then removed by centrifugation (12,000 x *g*, 30 min, 4 °C). The supernatant from this step was collected as the final cell extract and was stored in small aliquots and flash frozen for future use.

### Cell extract preparation [B]

An alternate CFPS expression reaction used lysate generated from *E. coli* Rosetta 2 (DE3) ΔlacZα^43^ (Table S3), largely as described above, except the culture was grown in a 100 L fermenter, and the lysate was treated by dialysis after lysis. This extract preparation was only used for PduA dilution assay (Fig. S1B).

### CFPS Reactions

CFPS was conducted according to methods previously described^25,42^. Briefly, 40 µL CFPS reactions were set up to produce proteins for analysis by confocal microscopy. The 40 µL CFPS reactions each contained: 10 mM Mg(Glu)_2_, 10 mM NH_4_(Glu), 130 mM K(Glu), 1.2 mM adenosine triphosphate (ATP), 0.85 mM guanosine triphosphate (GTP), 0.85 mM uridine 5’-triphosphate (UTP), 0.85 mM cytidine 5’-triphosphate (CTP), 0.034 mg/mL folinic acid, 0.171 mg/mL tRNA, 33.33 mM phosphoenol pyruvate (PEP), 2 mM 20 standard amino acids, 0.33 mM NAD+, 0.27 mM CoA, 4 mM Coenzyme-A (CoA), 1 mM putrescine, 1.5 mM spermidine, and 57 mM 4-(2-hydroxyethyl)-1-piperazineethanesulfonic acid (HEPES), 0.3 fraction extract, and a 1:100 dilution of DYKDDDDK Tag Monoclonal Antibody (FG4R), DyLight™ 550 (ThermoFisher Scientific Cat # MA1-91878-D550). To start the CFPS reaction, pJL1 plasmid (final concentration 13.33 ng/µL) containing our gene of interest was added to the reaction mixture. The CFPS reactions were pipetted into µ-Slide 18-well high glass bottom chambers (IBIDI Cat # 81817) and incubated at 30 °C for >2 h. ^14^C-leucine incorporation. The concentration of proteins produced in CFPS was quantified using ^14^C-leucine incorporation as previously described^24^. Briefly, 15 µL CFPS reactions included the 20 standard amino acids and 10 µM of ^14^C-leucine. CFPS reactions were incubated at 30 °C for two hours and then diluted 1:4 in 2 M urea and 1x Laemmli buffer (62.5 mM Tris pH 6.8, 2% sodium dodecyl sulfate (SDS), 10% glycerol, 0.05% bromophenol blue) with 10% β-mercaptoethanol. The diluted CFPS reactions were boiled at 95 °C for 20 minutes. Afterwards, 5 µL of the boiled CFPS reaction was added to 5 µL of 5 N KOH. Then, 4 µL of the KOH sample mix was spotted onto a glass fiber filter mat. The proteins were precipitated using 10% trichloroacetic acid. The filter mats were air dried and then covered in scintillation wax. Radioactive counts were measured from the precipitated proteins using liquid scintillation (PerkinElmer MicroBeta^2^ 2450 Microplate Counter) to get quantitative measurements of the protein yields produced in CFPS.

### Autoradiogram

In addition to using ^14^C-leucine incorporation, autoradiography was also used to verify protein production. The ^14^C-leucine incorporated CFPS reactions were boiled for 15 minutes at 95 °C and then run on an SDS-PAGE gel. The gel was with fixed with cellophane sheets soaked in deionized water and dried for one hour at 60 °C using a gel air dryer (Hoefer GD2001). The fixed gel was exposed to a phosphoscreen for 3-5 days. A Typhoon 7000 (GE Healthcare Life Sciences) was then used to scan the phosphoscreen using a 650 nm laser.

### Confocal/Super-resolution Microscopy

For imaging of 3D features of BMC-H assemblies, the 40 µL CFPS reaction was co-incubated with a 1:100 dilution of DYKDDDDK tag monoclonal antibody (FG4R), DyLight™ 550 (Invitrogen® Catalog No. MA1-91878-D550) in a µ-Slide 18 well glass bottom cell imaging chamber for examination with a Nikon SoRa spinning disk confocal microscope with an air 20x (0.8 NA) objective lens and a silicone-immersion 40x objective lens (1.25 NA). A Yokogawa CSU-W1 dual-disk spinning disk unit with 50μm pinholes, SoRa spinning disk (to increase magnification by 4x), and a Hamamatsu ORCA-Fusion Digital CMOS Camera were used for image acquisition. The microscope was controlled using Nikon imaging software NIS-Elements AR. All variants were imaged on at least 3 separate days. Microscope settings for each variant are described in Table S4. The SoRa microscope is housed in the Northwestern University Biological Imaging Facility supported by the NU Office for Research.

### Image Analysis

Maximum intensity projections were generated using ImageJ (version 1.54f) and cropped to known dimensions so scale bars could be added in Adobe Illustrator. 3D renderings were created using the Nikon NIS-Elements AR software and screenshots were taken to convert the images to 8-bit RGB images. Bound boxes were added in the Nikon software and then re-traced in Adobe Illustrator.

### PduA dilution assay

PduA was expressed from pJL1-PduA-FLAG plasmid added to the CFPS reaction at a concentration of 5 nM. 200 uL of CFPS reaction solution was placed per well of an eight-well chamber slide (80821, Ibidi, Gräfelfing, Germany) allowing the solution to cover the bottom of the well and was incubated at 37 °C overnight. A 20 uL sample from the 200 uL CFPS reaction was diluted into 100 uL water. 2 uL of the dilution was placed between a glass slide and coverslip for imaging. The experiment used a Zeiss Axio observer Z1 inverted microscope with an LD Plan-Neofluor 20x/0.4 Korr Ph2 M27 objective, operated in Brightfield mode. Images were captured with an Axiocam 506 camera using a 5 ms exposure time (Fig. S1A).

### Phenotype Quantification

All structures with quantified phenotypes were measured in FIJI. Hexagon diameters and fibril lengths were estimated on maximum intensity projections using the line tool from vertex to opposite vertex of a given hexagon (Fig S2A-B). Cluster diameters were measured for objects with relatively circular or spherical phenotypes. For these, maximum intensity projections were generated, and the draw tool was used to create regions of interest (ROIs) in a minimal area around the object in order to calculate the ferret diameter of the object (Fig. S2C). Hexagon thickness was estimated using the formula Z_object_ = | Z_slice 1_ – Z_slice 2_ | * (voxel height), where Z_object_ is the final calculated thickness of the object measured, Z_slice 1_ is the first Z slice wherein the hexagon appears, Z_slice 2_ is the final Z slice in which the object is in view, and voxel height is the distance between steps during acquisition on the microscope (Fig. S2D). This method overestimates thickness, as any object not perfectly parallel to the bottom of the microscope dish will appear in more slices, yielding a thicker measurement (Fig. S2E)

### Sequence alignments

All pairwise sequence alignments (Fig. 1B and Fig. 6A) were performed using EMBOSS Needle^44^. Green was added to highlight identical residues in Adobe Illustrator. Multiple sequence alignment (Fig. S4A) was performed using Clustal Omega version 1.2.4^44^.

### Protein Structure Prediction Using AlphaFold

The three-dimensional structure of CAETHG3289 was predicted using AlphaFold^35^. The input sequence was obtained from Stanley and Warren (2019) UniProt ID: UPI0003BAD469, and the prediction was performed using AlphaFold3^35^. Default parameters were used. Model confidence was assessed using the predicted Local Distance Difference Test (pLDDT) scores.

## Notes

### Competing Interest Statement

The authors have declared no competing interest.

