## Supplemental Figures and Tables for "Self-assembling protein materials with genetically programmable morphology and size"

**Supplemental Information**

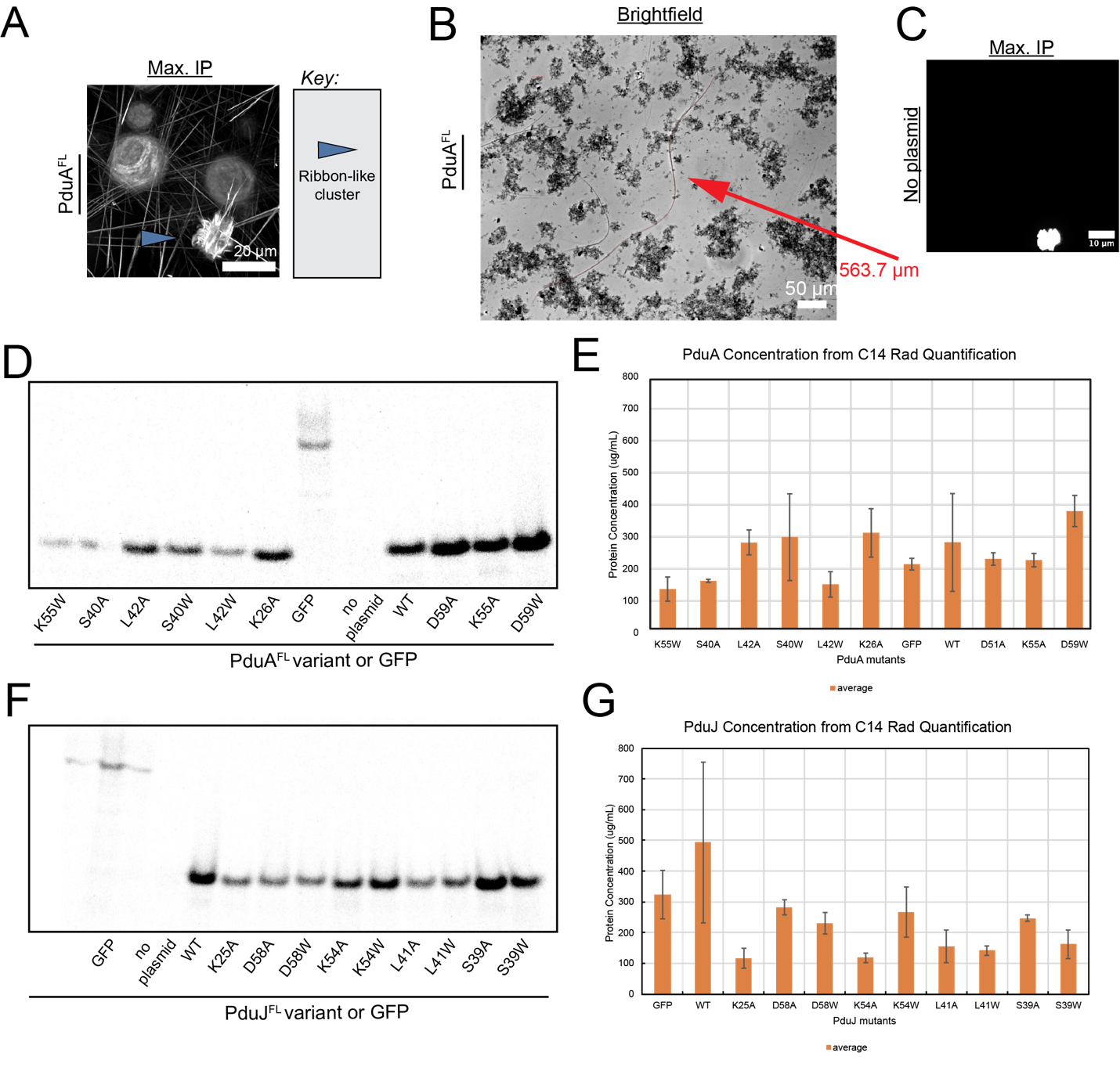

Figure S1: A) Maximum intensity projection of cell-free produced PduA. Blue arrow indicates ribbon-like cluster cited in-text. Scale bar = 20 µm. B) PduA dilution assay. Micrograph of cell-free produced PduA^FL^ diluted 1:5 and imaged using Brightfield. Red arrow highlights fibril measuring almost 0.6mm long. Scale bar = 50 µm. C) Maximum intensity projection of cell-free reaction with no plasmid added. Scale bar = 20 µm. D) Autoradiogram for GFP, no plasmid (controls), and all PduA^FL^ variants used in this study. E) Quantifications of protein expression of PduA^FL^ variants from C-14 Leucine incorporation in CFPS. F) Autoradiogram for GFP, no plasmid (controls), and all PduJ^FL^ variants used in this study. G) Quantifications of protein expression of PduJ^FL^ variants from C-14 Leucine incorporation in CFPS.

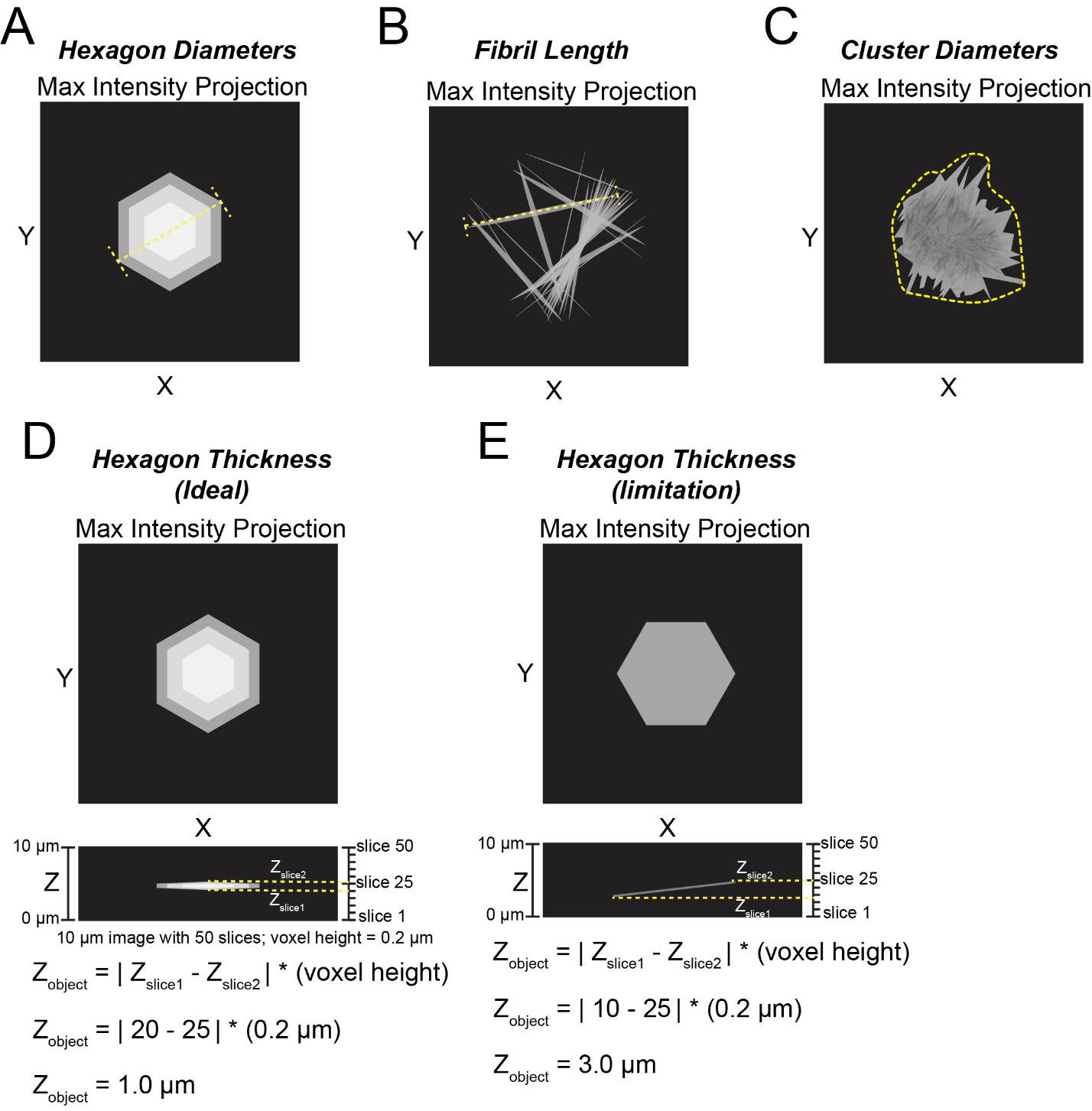

Figure S2. Schematic overview of phenotype quantification methods. A) Maximum hexagon diameters measured in FIJI using the line tool on the longest diameter of any hexagonal shaped object. B) Maximum fibril lengths measured in FIJI using the line tool. C) Clusters were defined as any low aspect-ratio objects that were not clearly hexagonal. Feret diameter was measured for clusters by drawing a minimal area shape around the object using the lasso tool in FIJI then measuring. D) Schematic overview of how object thickness was measured in an ideal scenario. E) Schematic overview demonstrating the limitation of using the method in (D) for measuring thickness of objects not perfectly parallel to the plane. Therefore, thickness was overestimated.

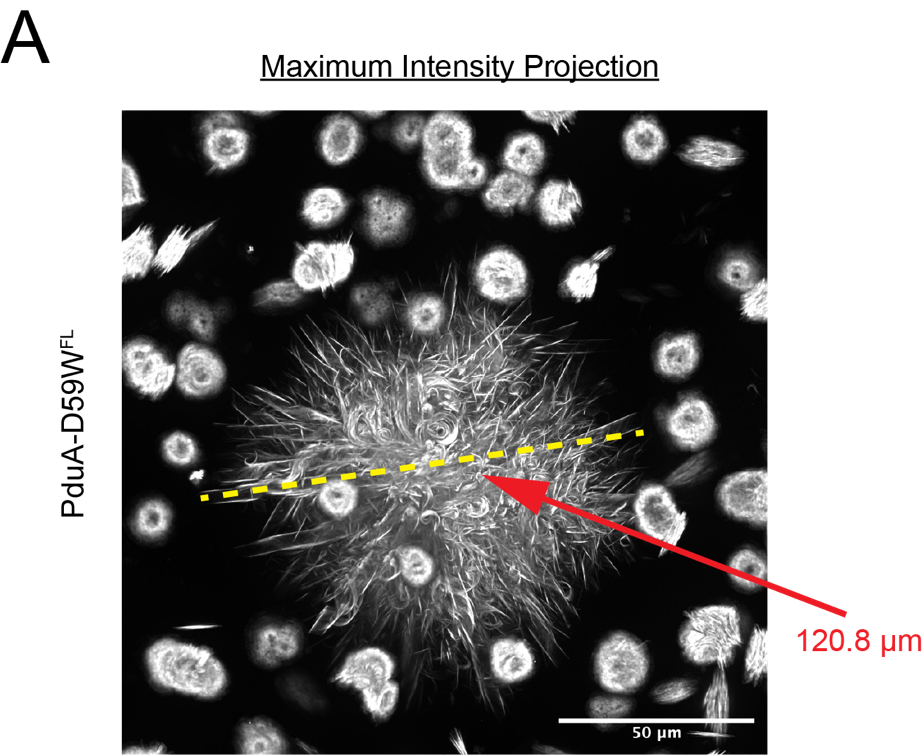

Figure S3: Maximum intensity projection of cell-free expressed PduA-D59W^FL^ demonstrating this variant assembles clusters of ribbons over 120 nm in diameter. Yellow dashed line indicates measured diameter and red arrow points to the structure in question, indicating it was measured at 120.8 nm (Fig. 3D). Scale bar = 50 µm).

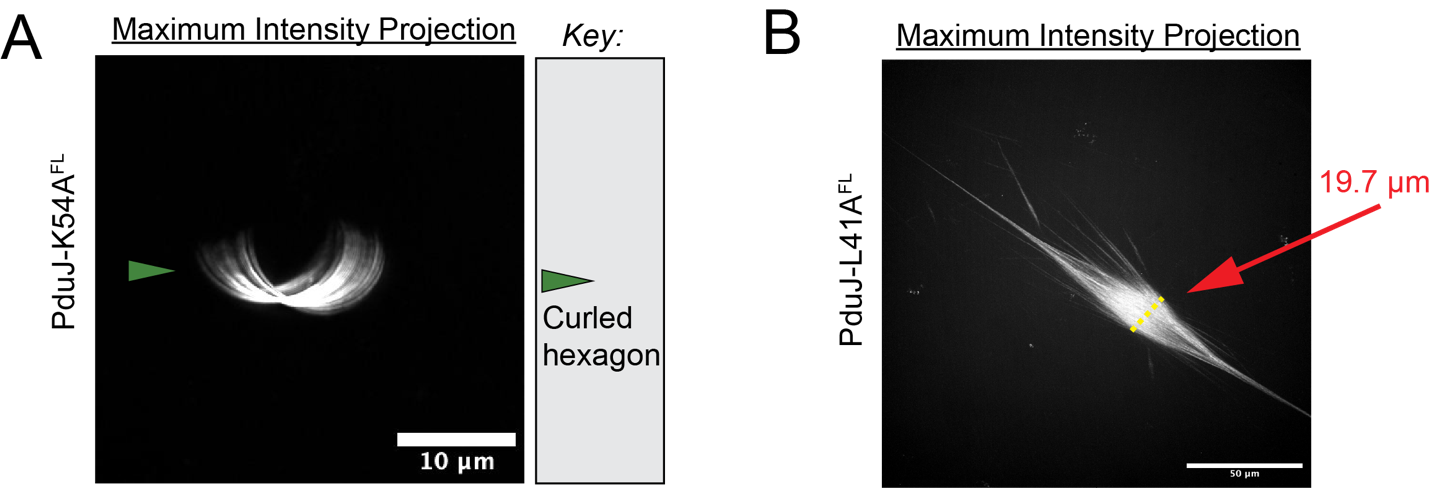

Figure S4: A) Maximum intensity projection of cell-free produced PduJ-K54A^FL^ demonstrating the curled hexagon morphology occasionally seen in PduJ-K54A^FL^. Green arrow points to curled hexagon structure. Scale bar = 10 µm. B) Maximum intensity projection of cell-free produced PduJ-L41A^FL^ demonstrating the diameter of the fibril bundles formed through the self-assembly process. Yellow line indicates the diameter and red arrow indicates the length measured 19.7 µm across. Scale bar = 50 µm.

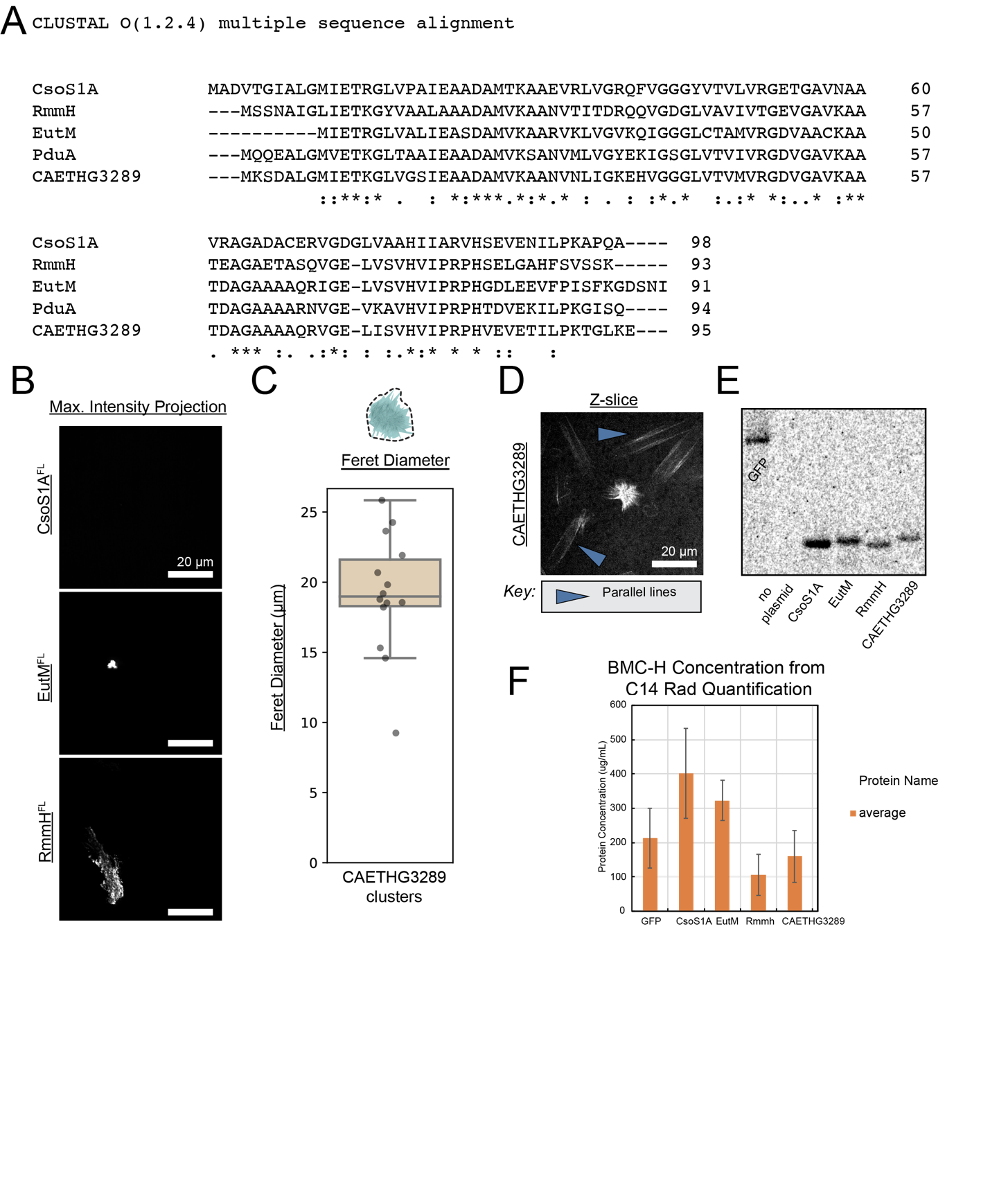

Figure S5: A) Multiple sequence alignment (MSA) of *E. coli* EutM, *M. smegmatis* RmmH, *H. neapolitanus* CsoS1A^,^ and *C. auto* CAETHG3289. MSA performed using Clustal O (1.2.4). B) Maximum intensity projections of cell-free produced CsoS1A^FL^, EutM^FL^, and RmmH^FL^. Scale bars = 20 µm. C) Box and whisker plot of diameter of CAETHG3289^FL^ clusters. D) Single z-slice from confocal micrograph of CAETHG3289^FL^ demonstrating phenotype of “two parallel lines” as mentioned in-text. Blue line highlights this phenotype. Scale bar = 20 µm. E) Autoradiogram for GFP, no plasmid (controls), and all BMC-H^FL^ variants used in this figure. F) Quantifications of protein expression of BMC-H^FL^ variants from C-14 Leucine incorporation in CFPS.

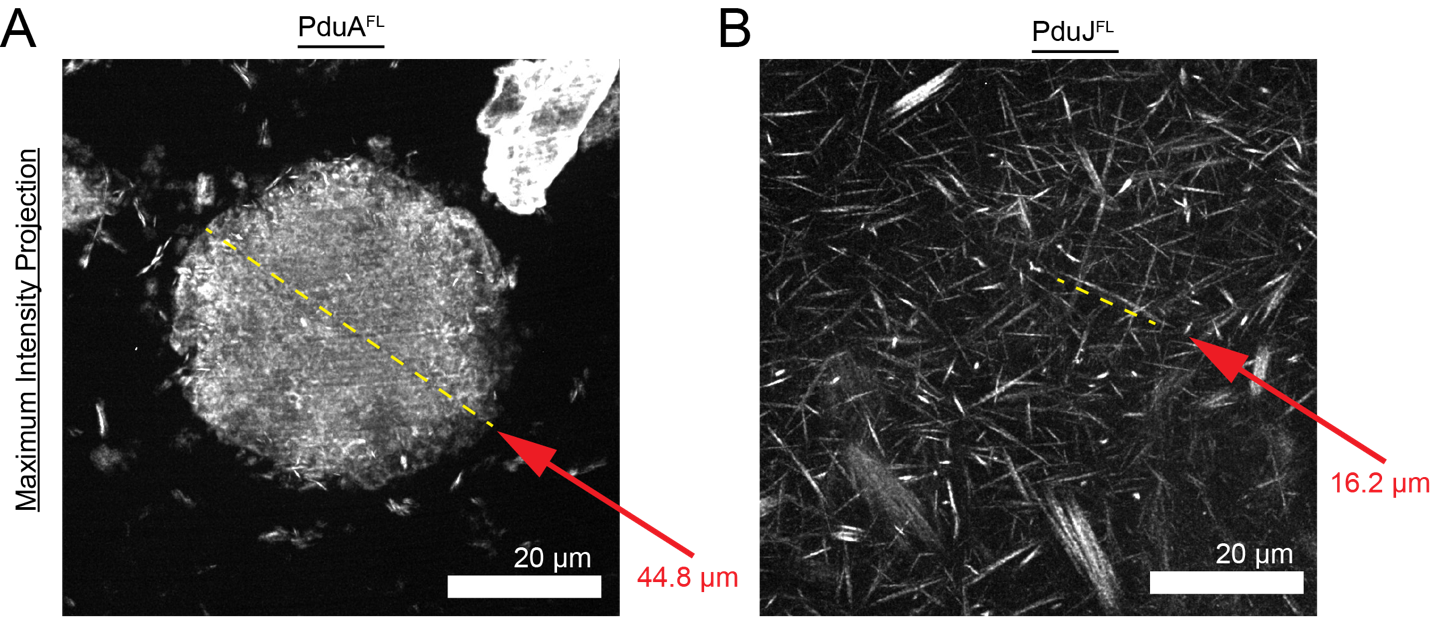

Figure S6. PduJ fibrils are shorter post-lyophilization and storage at 55ºC. A) Maximum intensity projection of cell-free produced PduA^FL^ that was lyophilized then immediately rehydrated in a water-antibody solution and imaged by super-resolution microscopy (Fig. 7A). Yellow dashed line demonstrates diameter of hexagon measured. Length measured in FIJI and marked in red to right of image. B) Maximum intensity projection of cell-free produced PduJ^FL^ that was lyophilized, stored at 55ºC for 1 week, rehydrated in a water-antibody solution, and imaged by super-resolution microscopy (Fig. 7A). Yellow dashed line demonstrates length of fibril measured. Length measured in FIJI and marked in red to right of image.

**Table S1.** Plasmids used in this study.

| **Name** | **Plasmid** | **Origin** | **Resistance** | **Plasmid Assembly Method** |
| --- | --- | --- | --- | --- |
| pCEM130 | pJL1-PduA-FLAG | colE1 | Kanamycin | Gibson Assembly |
| pCEM131 | pJL1-PduJ-FLAG | colE1 | Kanamycin | Gibson Assembly |
| CHAs196 | pJL1-PduA-K26A-FLAG | colE1 | Kanamycin | Gibson Assembly |
| CHAs197 | pJL1-PduJ-K25A-FLAG | colE1 | Kanamycin | Gibson Assembly |
| pCEM155 | pJL1-PduA-D59A-FLAG | colE1 | Kanamycin | Golden Gate Cloning |
| JBMs007 | pJL1-PduA-K55A-FLAG | colE1 | Kanamycin | QuikChange |
| JBMs012 | pJL1-PduA-D59W-FLAG | colE1 | Kanamycin | QuikChange |
| pCEM156 | pJL1-PduA-K55W-FLAG | colE1 | Kanamycin | Golden Gate Cloning |
| JBMs019 | pJL1-PduA-S40A-FLAG | colE1 | Kanamycin | Golden Gate Cloning |
| JBMs015 | pJL1-PduA-L42A-FLAG | colE1 | Kanamycin | QuikChange |
| JBMs020 | pJL1-PduA-S40W-FLAG | colE1 | Kanamycin | Golden Gate Cloning |
| pCEM154 | pJL1-PduA-L42W-FLAG | colE1 | Kanamycin | Golden Gate Cloning |
| JBMs013 | pJL1-PduJ-D58A-FLAG | colE1 | Kanamycin | Twist Bioscience |
| JBMs009 | pJL1-PduJ-K54A-FLAG | colE1 | Kanamycin | Twist Bioscience |
| JBMs014 | pJL1-PduJ-D58W-FLAG | colE1 | Kanamycin | Twist Bioscience |
| JBMs010 | pJL1-PduJ-K54W-FLAG | colE1 | Kanamycin | Twist Bioscience |
| JBMs021 | pJL1-PduJ-S39A-FLAG | colE1 | Kanamycin | Twist Bioscience |
| JBMs017 | pJL1-PduJ-L41A-FLAG | colE1 | Kanamycin | Twist Bioscience |
| JBMs022 | pJL1-PduJ-S39W-FLAG | colE1 | Kanamycin | Twist Bioscience |
| JBMs018 | pJL1-PduJ-L41W-FLAG | colE1 | Kanamycin | Twist Bioscience |
| pAJA009 | pJL1-CAETHG-FLAG | colE1 | Kanamycin | Twist Bioscience |
| CHAs194 | pJL1-CsoS1A | colE1 | Kanamycin | Gibson Assembly |
| CHAs193 | pJL1-EutM-FLAG | colE1 | Kanamycin | Gibson Assembly |
| CHAs195 | pJL1-RmmH-FLAG | colE1 | Kanamycin | Gibson Assembly |
| pCHA001 | pJL1-sfGFP | colE1 | Kanamycin | gift from Jewett lab |

**Table S2.** Primers used in this study.

| **Name** | **Purpose** | **Description** | **Sequence** |
| --- | --- | --- | --- |
| CHA17 | Gibson Assembly | Amplify pduA with GA overhang For | ttaagaaggagatatacataATGCAACAAGAAGCACTAG |
| CHA18 | Gibson Assembly | Amplify FLAG tag and GA overhang Rev | tttgttagcagccggtcgacTTACTTGTCATCGTCATCTTTATAATC |
| CHA20 | Gibson Assembly | Amplify pduJ with GA overhang For | ttaagaaggagatatacataATGAATAACGCACTGGGAC |
| NWKo483 | Golden Gate cloning | Amplify *pduA* with GG overhang For | aGGTCTCaCATGCAACAAGAAGCACTAGGAATG |
| NWKo494 | Golden Gate Cloning | Amplify *pduA* with FLAG tag and GG overhang Rev | tGGTCTCatTTACTTGTCATCGTCATCTTTATAATCttgg |
| NWKo422 | Golden Gate Cloning | K55W | aGGTCTCAcggcgatgttggcgcggtcTGGgcggccaccgatgcaggtgccgcagccgcacgcaacgtgggtgaagtgaaagccgtacacgtcatcccactGAGAC |
| NWKo455 | Golden Gate Cloning | L42W | aGGTCTCAaatggttaagtcagccaatgtgatgttagtgggctatgaaaagattggctccgggTGGgtaaccgtcatcgtgcgcggcgatgttggcgcggtGAGAC |
| NWKo425 | Golden Gate Cloning | D59A | aGGTCTCAcggcgatgttggcgcggtcaaagcggccaccGCGgcaggtgccgcagccgcacgcaacgtgggtgaagtgaaagccgtacacgtcatcccactGAGAC |
| JBMo001 | QuikChange | *pduA-L42A* For | ggctccgggGCggtaaccgtcatcg |
| JBMo002 | QuikChange | *pduA-L42A* Rev | gatgacggttaccGCcccggagccaatc |
| JBMo003 | QuikChange | *pduA-K55A* For | GttggcgcggtcGCGgcggccacc |
| JBMo004 | Quikchange | *pduA-K55A* Rev | CggtggccgcCGCgaccgcgcc |
| JBMo005 | QuikChange | *pduA-D59W* For | gcggccaccTGGgcaggtgccg |
| JBMo006 | QuikChange | *pduA-D59W* Rev | ctgcggcacctgcCCAggtggcc |
| NWKo484 | Golden Gate cloning | Amplify *pduJ* with GG overhang For | aGGTCTCaCAtgaataacgcactgggactgg |
| NWKo486 | Golden Gate cloning | Amplify *pduJ* with FLAG tag and GG overhang Rev | tGGTCTCatTTACTTGTCATCGTCATCTTTATAATCggc |
| CHAo256 | Gibson Assembly | Amplify *csoS1A* with GG overhang For | ttaagaaggagatatacataATGGCTGATGTAACTGGTATTGCTCTGG |
| CHAo255 | Gibson Assembly | Amplify *eutM* with GG overhang For | ttaagaaggagatatacataATGATTGAGACTAGAGGTTTGGTTGCGT |
| CHAo257 | Gibson Assembly | Amplify *rmmH* with Gibson overhang For | ttaagaaggagatatacataATGAGCAGTAATGCAATCGGCCTTATTG |

**Table S3.** Strains used in this study.

| **Strain** | **Organism** | **Genotype** |
| --- | --- | --- |
| CEMs034 | *E. coli* BL21 (DE3) | Wild type |
| *N/A* | *E. coli* Rosetta 2(DE3) | ΔlacZα |

**Table S4.** Microscope settings used for different variants throughout the study. All settings below used excitation laser of 561 nm and SoRa magnification of 4X using a 50 µm pinhole. Images were all deconvolved with type-blind, 20 iterations. ^a^Fig. 1B, first image. ^b^Fig. 1B, second image & Fig. 1C. ^c^Fig. 5A, first image. ^d^Fig. 5A, second image & Fig. 5C.

| **Variant Name** | **Exposure Time (ms)** | **Laser Power (%)** | **Objective Lens (X)** |
| --- | --- | --- | --- |
| PduA^FL^ | 200 | 25^a^, 50^b^ | 20 |
| PduA-K26A^FL^ | 200 | 50 | 40 |
| PduJ^FL^ | 200 | 50 | 20 |
| PduJ-K25A^FL^ | 500 | 40 | 40 |
| PduA-D59A^FL^ | 200 | 25 | 20 |
| PduA-K55A^FL^ | 200 | 50 | 20 |
| PduA-D59W^FL^ | 200 | 50 | 20 |
| PduA-K55W^FL^ | 500 | 40 | 40 |
| PduA-S40A^FL^ | 200 | 50 | 20 |
| PduA-L42A^FL^ | 200 | 50 | 20 |
| PduA-S40W^FL^ | 200 | 50 | 20 |
| PduA-L42W^FL^ | 500 | 40 | 40 |
| PduJ-D58A^FL^ | 400 | 50 | 40 |
| PduJ-K54A^FL^ | 400 | 50^c^, 75^d^ | 20 |
| PduJ-D58W^FL^ | 200 | 40 | 20 |
| PduJ-K54WFL | 400 | 50 | 40 |
| PduJ-S39A^FL^ | 200 | 50 | 20 |
| PduJ-L41A^FL^ | 200 | 40 | 20 |
| PduJ-S39W^FL^ | 200 | 20 | 20 |
| PduJ-L41W^FL^ | 200 | 25 | 20 |
| CAETHG3289^FL^ | 400 | 50 | 20 |
| CsoS1A^FL^ | 200 | 75 | 20 |
| EutM^FL^ | 200 | 75 | 20 |
| RmmH^FL^ | 200 | 75 | 20 |
